# Structural implications of BK polyomavirus sequence variations in the major viral capsid protein Vp1 and large T-antigen: a computational study

**DOI:** 10.1101/2023.12.21.572635

**Authors:** Janani Durairaj, Océane M. Follonier, Karoline Leuzinger, Leila T Alexander, Maud Wilhelm, Joana Pereira, Caroline A. Hillenbrand, Fabian H. Weissbach, Torsten Schwede, Hans H. Hirsch

## Abstract

BK polyomavirus (BKPyV) is a double-stranded DNA virus causing nephropathy, hemorrhagic cystitis, and urothelial cancer in transplant patients. The BKPyV-encoded capsid protein Vp1 and large T-antigen (LTag) are key targets of neutralizing antibodies and cytotoxic T-cells, respectively. Our single-center data suggested that variability in Vp1 and LTag may contribute to failing BKPyV-specific immune control, and impact vaccine design. We therefore analyzed all available entries in GenBank (1516 *VP1*; 742 *LTAG)* and explored potential structural effects using computational approaches. BKPyV-genotype (gt)1 was found in 71.18% of entries, followed by BKPyV-gt4 (19.26%), BKPyV-gt2 (8.11%) and BKPyV-gt3 (1.45%), but rates differed according to country and specimen type. Vp1-mutations matched a serotype different than the assigned one or were serotype-independent in 43%, 18% affected more than one amino acid. Notable Vp1-mutations altered antibody-binding domains, interactions with sialic acid receptors, or were predicted to change conformation. LTag-sequences were more conserved, with only 16 mutations detectable in more than one entry and without significant effects on LTag-structure or interaction domains. However, LTag changes were predicted to affect HLA-class I presentation of immunodominant 9mers to cytotoxic T-cells. These global data strengthen single center observations and specifically our earlier findings revealing mutant 9mer epitopes conferring immune escape from HLA-I cytotoxic T cells. We conclude that variability of BKPyV-Vp1 and LTag may have important implications for diagnostic assays assessing BKPyV-specific immune control and for vaccine design.

**IMPORTANCE:** Type and rate of amino acid variations in BKPyV may provide important insights into BKPyV diversity in human populations and an important step towards defining determinants of BKPyV-specific immunity needed to protect vulnerable patients from BKPyV diseases. Our analysis of BKPyV sequences obtained from human specimens reveals an unexpectedly high genetic variability for this double-stranded DNA virus that strongly relies on host cell DNA replication machinery with its proof reading and error correction mechanisms. BKPyV variability and immune escape should be taken into account when designing further approaches to antivirals, monoclonal antibodies and vaccines for patients at risk of BKPyV diseases.

## INTRODUCTION

BK polyomavirus (BKPyV) is one of more than 10 human polyomaviruses (HPyVs) which belong to the *polyomaviridae* family found in nearly all vertebrates (1, 2). BKPyV infects more than 90% of the general human population without specific signs or symptoms but can cause significant diseases in immunocompromised patients (3–5). The leading entities are nephropathy, and hemorrhagic cystitis, which complicate 5% to 25% of mostly kidney and allogeneic hematopoietic cell transplant recipients, followed by approximately 0.1% of urothelial cancer cancers carrying a chromosomal integration of BKPyV (6, 7). Pathology, rate and risk factors of BKPyV diseases differ in different patient populations, but their underlying common theme is insufficient adaptive immunity to BKPyV (8). Since there are no effective antiviral drugs for treatment or prevention of BKPyV replication and associated diseases (9), current clinical management relies on reconstituting BKPyV-specific humoral and cellular immunity. However, our earlier work has identified significant changes in 9mer epitopes which were associated with failure of CD8 T cells to activate polyfunctional responses, to kill and to proliferate (10–12). Similarly, escape from neutralizing antibodies is suspected of contributing to escape from humoral adaptive immune control (13–16).

BKPyV virions are non-enveloped icosahedral capsids of 40-45 nm diameter formed by 72 pentamers of the major capsid protein Vp1 outside and one Vp2 and one Vp3 inside at a ratio of 5:1:1. Inside, the circular double-stranded DNA genome of approximately 5.1 kb is packaged using host cell-derived histones (8). Akin to other HPyVs, the BKPyV genome can be divided into three major regions called the non-coding control region (*NCCR*), the early viral gene region (*EVGR*) and the late viral gene region (*LVGR*). The *NCCR* harbors the origin of viral DNA replication and bidirectional intertwined promoter/enhancer sequences. Together with host cell factors, the *NCCR* regulates the sequential expression of the *EVGR*-encoded regulatory large and small T-antigen (LTag, sTag), the viral DNA replication and expression of the *LVGR-* encoded regulatory agnoprotein, and the structural capsid proteins Vp1, Vp2, and Vp3 (17–19). Further on the *LVGR-*strand, two micro-RNAs are found downstream of the *VP1-* polyadenylation signal and which downregulate *LTAG*-transcripts as well as *ULBP3*-transcripts, a potential target of natural killer-lymphocytes (20).

The *VP1*-sequence variability of circulating BKPyV gives rise to four major Vp1 serotypes, initially defined by neutralizing antibody (NAb) titers (21, 22). More recent phylogenetic analyses use larger genome sequences that also include parts of the *EVGR* and currently define 12 BKPyV subgroups (1). Humoral immunity to the intracellular LTag and sTag cannot confer protection by NAbs. However, cellular immunity to these *EVGR*-encoded proteins seems to play a critical role which involves immunodominant 9mer peptide clusters presented by HLA-class I molecules to cytotoxic CD8 T lymphocytes (CTLs) (23–25). Notably, genotype-dependent and genotype-independent variability in the LTag-sequence has been linked to reduced 9mer-directed CTL responses (10, 11). Thus, sero- and genotype-encoded variability in Vp1 and LTag may impair BKPyV-specific immune control by NAbs and cytotoxic T cells in transplant patients, respectively.

Given the potential impact on diagnostic assays and vaccine design, we set out to analyze the variability of the BKPyV *VP1* and *LTAG* protein coding sequences from public databases and from our recent molecular study on hematopoietic cell transplantation (HCT) recipients (10) using computational approaches. Our findings revealed that 43% of *VP1* sequences had non-synonymous changes, whereby mutations in 23 amino acid positions were highly prevalent. We analyzed the potential effects on Vp1 structure, especially those interacting with cellular sialic acid receptors, NAbs, and the minor capsid proteins Vp2 and Vp3. Additionally, we explored changes within the LTag protein and assessed their effect on confirmed immunodominant CTL epitopes including their cross-protective potential.

## MATERIALS & METHODS

### Mapping variants from GenBank entries

Nucleic acid sequences were denoted in capital letters in *italics*, while the encoded respective proteins used at least one small letter and no italics (8). We retrieved all available entries for BKPyV *LTAG* nucleic acids and encoded LTag amino acid sequences as well as *VP1* nucleic acids and encoded Vp1 protein sequences (NCBI Taxonomy IDs: 1891762, 1417981, 1303334, 10631), for JC polyomavirus (JCPyV) (NCBI Taxonomy ID: 10632) and for simian virus 40 (SV40) (NCBI Taxonomy ID: 1891767) from GenBank (26) as of May 2023 using Entrez tblastn search (27) with the protein sequences of AB211371 for BKPyV, NC_001699 for JCPyV, and AF155358 for SV40. This was followed by an Entrez free text search using “VP1” or “LTAG” as the query and restricted to the respective NCBI taxonomy IDs for each of the three polyomaviruses. The GenBank identifiers from both search strategies were combined and Vp1 and LTag amino acid sequences were extracted from the translated CDS defined in the corresponding GenBank files. If no CDS containing “VP1”, “VP-1”, “major capsid”, “viral protein-1” for Vp1 or “large” for LTag was found, sequences were extracted from the translated tblastn results. Information about the sample source and country of origin were obtained from the corresponding GFF files for each entry. The resulting protein sequences were aligned using MUSCLE (version 5.1) (28) using default settings. All mutation statistics reported in the text are based on the positions in this alignment which are present in the BKPyV Vp1 and LTag sequences of AB211371. The Vp1 alignment and LTag alignment files are provided as Supplementary Files 1 and 2 respectively.

### Genotyping of BKPyV Vp1 and LTag sequences

BKPyV Vp1 and LTag sequences were grouped using the BKPyV reference sequences (**Table 1**). The Vp1 reference sequences were previously identified to correspond to four different serotypes (21, 22), while the LTag reference sequences define the LTag genotype. The closest reference sequence was found for each entry (Vp1 or LTag) sequence, defined as the sequence with the lowest number of amino acid changes from the entry within the positions which differed in at least one reference sequence (these are referred to as serotype-defining positions for Vp1, SDP). Each entry was assigned to the Vp1 serotype or LTag genotype of its closest reference sequence. An amino acid position in a viral sequence was defined as a mutation if the encoded amino acid does not occur in any of the reference sequences of the assigned serotype/genotype. Mutations resulting in amino acids present in a different serotype than the one assigned were referred to as serotype-exchange mutants (SXM), and all other mutations were denoted serotype-independent mutants (SIM). An amino acid position in a BKPyV viral sequence was defined as being different from JCPyV or SV40 if all the amino acids found in the GenBank JCPyV or SV40 entries differed from the amino acids found in any of the BKPyV reference sequences at that position. For each *VP1* nucleotide sequence, mutations from the corresponding reference nucleotide sequence on the antisense DNA strand were recorded if they matched the APOBEC3 mutational signature, defined as C->G and C->T mutations in a T<u>C</u>T or T<u>C</u>A context.

**Table 1.**
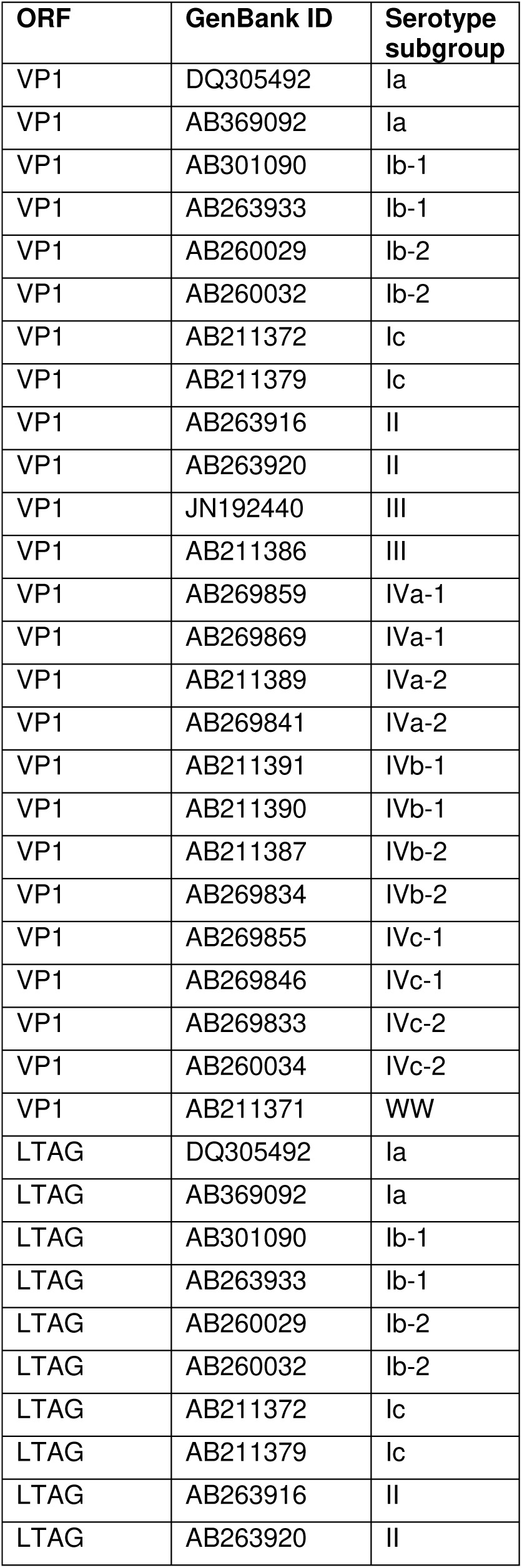
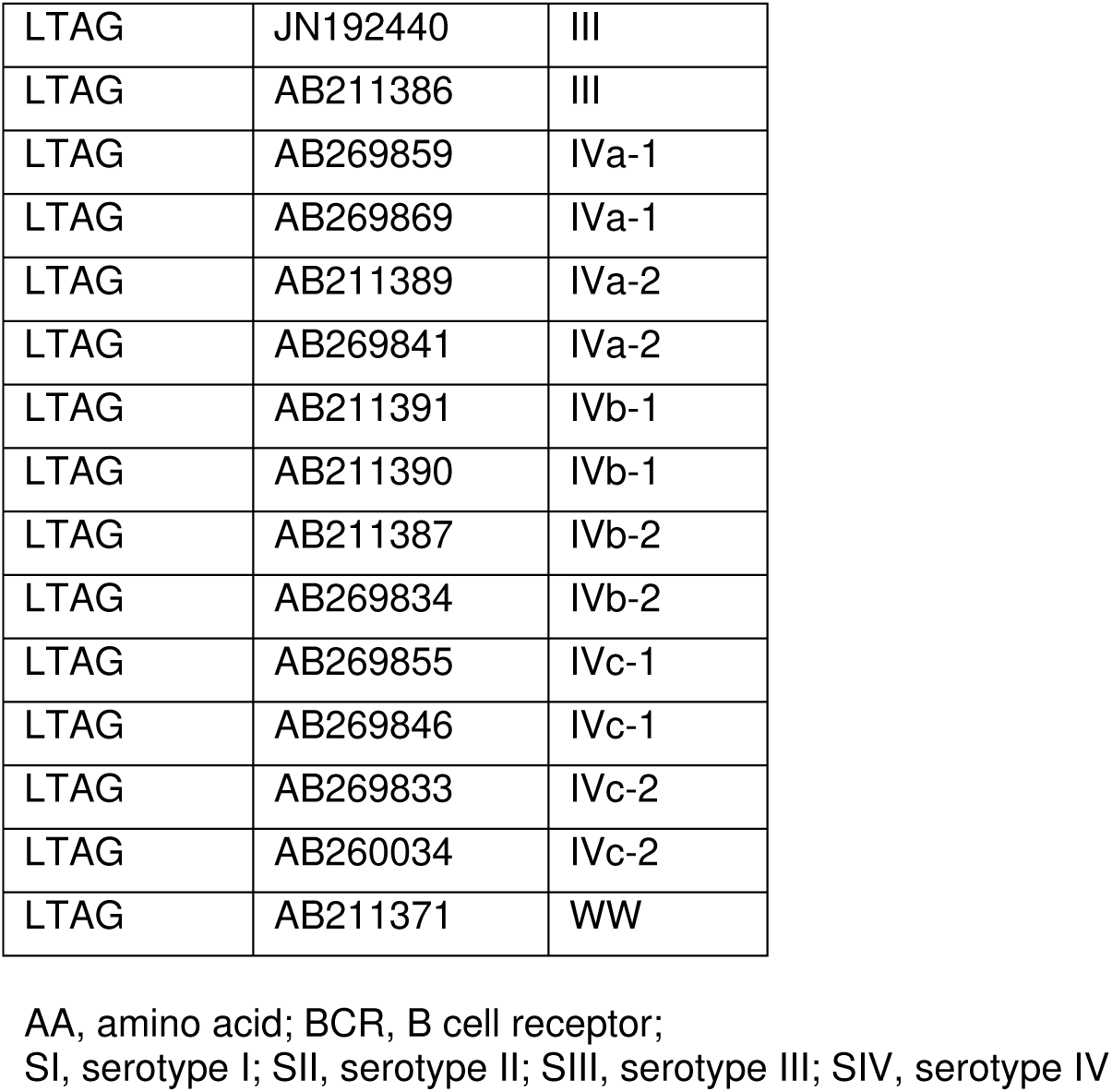
Reference sequences for BKPyV Vp1 and LTag.

### Vp1 experimental structure analysis

To obtain an overview of the conformational flexibility within the Vp1 pentamer, all chains of 9 Vp1 structures of BKPyV (PDB IDs: 7ZIQ, 6GG0, 6ESB, 7B6C, 7B6A, 4MJ1, 5FUA, 7B69, 4MJ0) were superposed (**Supplementary Figure 1**). Since this superposition appeared to have an unexpected structural conservation of the BC-loop, the per-residue median electron density support for individual atoms (EDIA_m_) scores (29) were obtained and averaged across 3 Vp1 crystal structures (PDB IDs: 4MJ0, 4MJ1 and 7ZIQ). Using a median EDIA_m_ < 0.8, allowed to mark residues where the electron density data has a poor fit to the coordinates in the crystal structure (29).

### Vp1 structural modelling

The Vp1 pentameric model was obtained using the SWISS-MODEL automated modelling workflow (30), (31). The per-residue Quantitative Model Energy Analysis (QMEAN) (32) structure quality metric for this model was used to narrow focus to residues predicted with high confidence (>0.65 QMEAN). Sialic acid receptors (from PDB IDs: 6ESB, 4MJ0), antibodies (from PDB IDs: 6GG0, 7PA7), and the Vp2/Vp3 binding fragment (from PDB ID: 1CN3) were transplanted to the Vp1 model using the PyMol super command (33) . Residues participating in interactions were obtained by considering atoms within 6 Å from the interacting partner and passing the QMEAN quality threshold of 0.65.

### Vp1 epitope prediction

DiscoTope 3.0 (version 1.1a) (34) was used to predict neutralizing antibody epitopes for each individual chain of the modelled Vp1 structure. Residues passing the default DiscoTope prediction threshold of -7.7 in at least 2 chains and passing the QMEAN quality threshold of 0.65 were considered as predicted BCR epitope residues. NetMHCPan (version 4.1) (35) was used to predict CD8 T-cell epitopes for the most common HLA-A, -B, and -C types. Only predicted strong binders (based on a rank threshold of 0.5) were considered.

### Vp1-antibody interaction mutants

Prediction of mutations likely to be involved in immune escape were obtained using FoldX (36) mutagenesis of PDB 7PA7. The BuildModel command was used to build mutant models with 3 replicates for 30 point-mutations both with (holo) and without (apo) the bound antibody. The differences in predicted ΔΔG between all pairs of apo-holo models across replicates was used to hypothesize which mutations may destabilize the antibody-binding interface. Hydrogen bonds between Vp1 atoms and antibody atoms are highlighted using PLIP (37) .

### LTag structural modelling and annotation

A structural model was predicted for LTag using AlphaFold (38) with default settings, and split into domains (defined as in (10)). All structure interaction annotations are transferred from SV40 LTag structures. This includes intra-hexamer interactions for the Helicase domain obtained as those residues within 6Å of another Helicase chain in all SV40 LTag crystal structures (PDB IDs: 1SVO, 1N25, 1SVL and 1SVM), and residues binding to p53 (39), ATP (40), zinc (41), and DNA (42). Confirmed immunodominant 9mer epitopes presented by HLA-class I to CD8 T cells and their frequencies were obtained as observed in the checkerboard analysis of kidney transplant patients in the Swiss Transplant Cohort Study (10). Matches of the 9mer epitopes to the human proteome were obtained using PepMatch (43) with a *k*-mer size of 5. NetMHCPan (version 4.1) (35) was used to predict CD8 T-cell epitopes for the 9 most common each of HLA-A, -B, and -C types. Only predicted strong binders based on a rank threshold of 0.5 were considered.

## RESULTS

### Diversity of the BKPyV capsid protein Vp1

#### Geographic origin and serotype variation of the BKPyV-Vp1

We analyzed all available BKPyV *VP1* sequences in GenBank to evaluate variation as a proxy of variability among sequences from diverse sources. Our dataset comprised 1516 sequence entries, of which 650 displayed variations from serotype reference sequences, resulting in 288 unique combinations of mutations. The distribution of Vp1 serotypes confirmed BKPyV serotype-I as being most common (71.2% of entries), followed by serotype-IV (19.3%) and -II (8.1%) with some variation according to the geographical region (**Figure 1A**). BKPyV serotype-III was rare (<2%) and mostly derived in Japan, Spain, or USA. The majority of sequences were derived from urine samples, followed by blood and organ specimens (**Figure 1B**). Overall, no significant association between serotype, source of specimen and number of mutations was observed (**Figure 1B and 1C**). However, a study from Brazil reported mostly BKPyV serotype-II in saliva samples (44). 43% of Vp1 sequences exhibited deviation from the assigned serotype, and 18% had three or more amino acid mutations. Of 362 amino acids in Vp1, 185 (51% of the protein) were mutated in at least one entry compared to the assigned serotype; 65 positions (18%) were supported by at least three entries and most of them occurred in serotype-I (**Figure 1C**). Taken together, these findings demonstrate an overall dominance of serotype-I and -IV among the reported BKPyV Vp1 entries, while pointing to geography and type of specimen potentially contributing to differences in BKPyV-serotype prevalence and variability.

**Figure 1.**
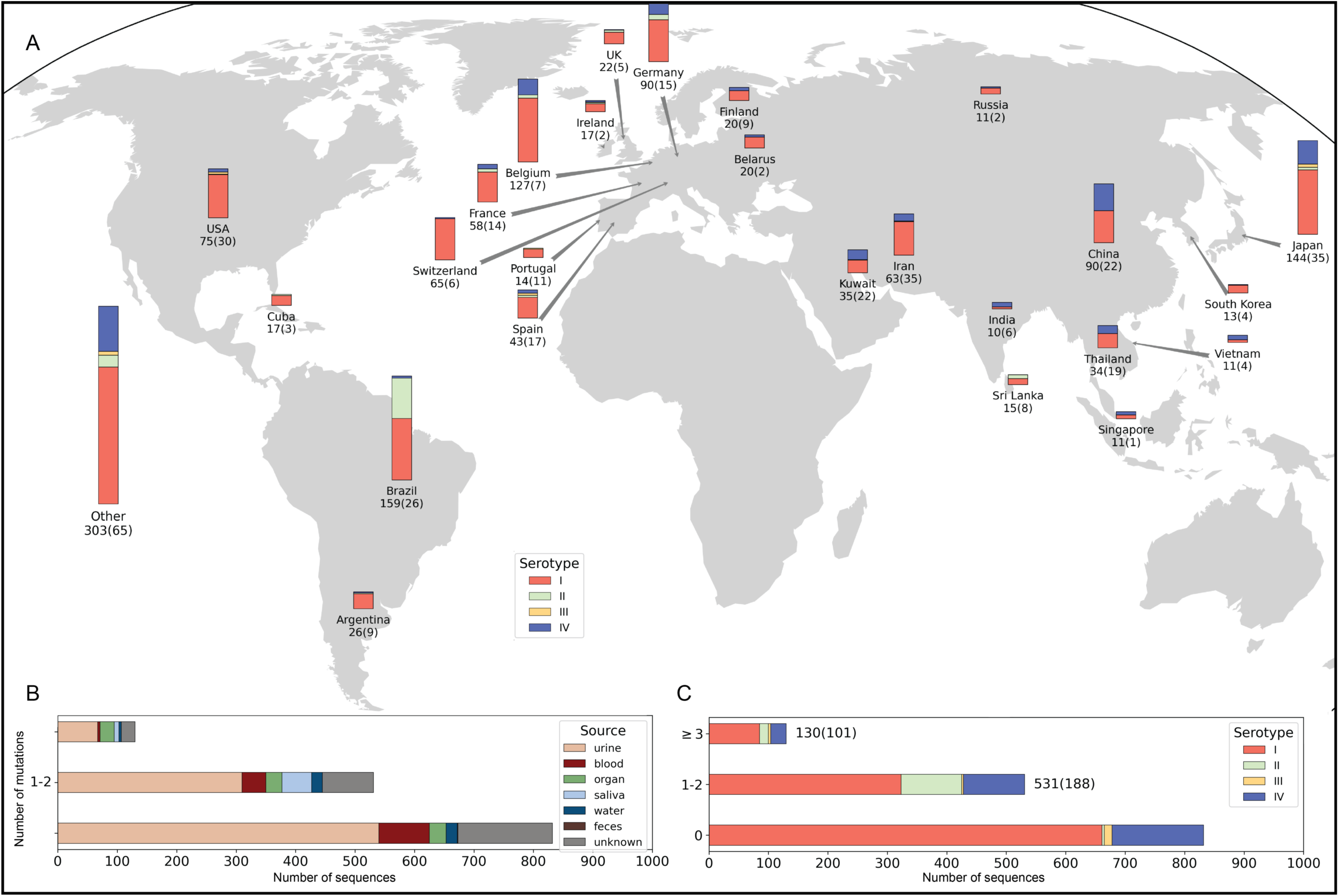
BKPyV serotype and variability across countries and specimen type. A. Vp1 sequence and serotype according to geographical source. The total number of sequences and number of unique sequences (in brackets) for each country are provided below the country name. Countries with fewer than 10 sequences or missing country data in GenBank metadata are combined under “Other”. B. Number of Vp1 amino acid changes according to specimen source and number of entries. C. Number of Vp1 amino acid changes according to serotype and number of entries.

To investigate whether variability concerned particular parts of the 362 amino acid-long BKPyV-Vp1 protein, we visualized amino acid changes in the linear sequences (**Figure 2**). The results identified mutation hot-spots in the regions around amino acids 50 – 90, 110 – 130 and 160 – 180 (**Figure 2, center panel**). From the set of 17 BKPyV-Vp1 reference sequences (**Table 1**), 29 amino acid positions differed in at least one sequence and were denoted as serotype-defining positions (SDPs). For 452 mutations in 24 positions (6.6% of the Vp1 protein), the variant amino acid occurred in a SDP and corresponded to a serotype-exchange mutation (SXM), since the altered amino acid belonged to a different serotype than the one assigned to the overall sequence. For 834 mutations in 178 positions (49.2% of the Vp1 protein), the amino acid corresponded to a serotype-independent mutation (SIM), as it did not correspond to any serotype-defining or reference amino acid.

**Figure 2.**
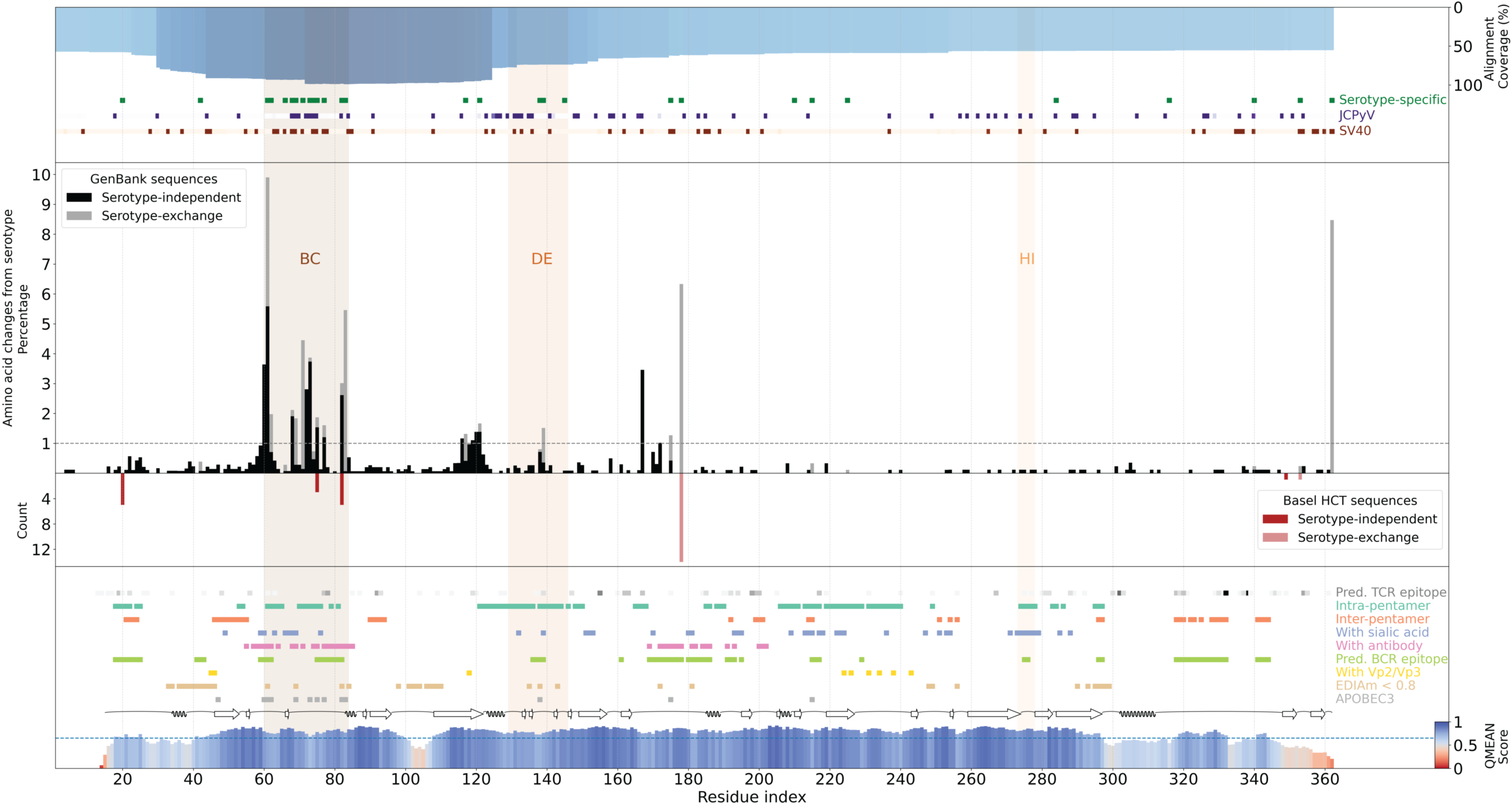
BKPyV Vp1 sequence mutations and annotations. Percentage of serotype-independent mutations (black bars) and serotype-exchange mutations (grey bars) at each Vp1 residue position of the GenBank entries. The grey dashed line represents 1%, above which variants are discussed further. Shaded areas mark residues within the Vp1 BC-, DE- and HI-loops. Mutants identified in patients of a recent study in Basel are shown as counts of serotype-independent (red bars) and serotype-exchange mutations (light red bars). The top blue bar indicates alignment coverage at each position. Green squares mark residues with serotype-specific changes BKPyV Vp1. Purple and orange squares mark residues where none of the reference serotype amino acids are found in corresponding positions in JCPyV and SV40 polyomavirus Vp1, respectively. The Vp1 structure coverage and quality is shown on a red-white-blue scale on the bottom. Colored squares mark residues involved in intra-pentamer contacts, inter-pentamer contacts, contacts with sialic acid receptors (from PDB: 6ESB, 4MJ0), contacts with antibodies (from PDB: 6GG0, 7PA7), predicted B-cell epitope residues, contacts with Vp2/Vp3 (from PDB: 1CN3), residues with lower electron density fit across available crystal structures (avg. EDIAm <0.8), and residues with APOBEC3-like mutations detected in at least two sequences. Contacts are only considered for residues with structure quality (QMEAN) over 65 (blue dashed line).

The alignment coverage was highest with >90% for the region from the BC-loop to the DE-loop and gradually decreased towards 60% at the N-terminus and C-terminus, respectively (**Figure 2**, **top panel**). SDPs of BKPyV-Vp1 were found over the entire length of the protein, but clearly clustered around the BC-loop and DE-loop. A similar clustering was also observed for JCPyV and SV40 Vp1 amino acids distinct from BKPyV-Vp1 (**Figure 2**, **top panel**, *dot plots, green versus purple and red*). Of note, the BC-loop region showed the highest rate of SXM and SIM (**Figure 2**, **center panel**, *grey and black bars*). Residues in region (110–130) preceding the DE-loop also exhibited considerable variation within BKPyV-Vp1 (**Figure 2**, **center panel**).

To explore the potential impact of the mutations on structure, function and immunogenicity, we used a homology model (constructed with SWISS-MODEL) of the chain-1 through -5 of one Vp1 pentamer and included one Vp1 (chain-1’) of a neighboring pentamer. On the linear Vp1 sequence, we marked intra-pentameric interactions, inter-pentameric interactions, and interactions with other relevant entities (**Figure 2, bottom panel**). To identify high confidence interacting amino acid residues, we estimated the quality of the predicted structure by obtaining the *per* residue average QMEAN score of the combined Vp1 chain-1 to -5 together with chain 1’ (**Figure 2, bottom panel**, red-blue scale). To provide an overview of amino acid residues potentially affecting the humoral and cellular adaptive immune response, we also included predictions of CTL epitope presentation to the 9 most common alleles each of HLA-A, B and C, as well as predictions of the neutralizing antibody epitopes (**Figure 2, bottom panel,** dot plots). Several positions showed changes matching the profile of the human Apolipoprotein B editing complex 3 (APOBEC3) mutational signatures (**Figure 2, bottom panel**, gray dots; see also **Table 2**).

**Table 2.** APOBEC3-like mutation signatures BKPyV *VP1*. Amino acid residue number, serotype, amino acid change, and trinucleotide change of mutations matching the APOBEC3 signature.

| **Vp1 AA residue** | **Serotype: AA mutation (counts)** | **Mutation (trinucleotide)** |
| --- | --- | --- |
| 6 | SI: R6K (1) | TCT -> TTT |
| 47 | SI: E47Q (2) | TCT -> TGT |
|  | SI: E47K (1) | TCT -> TTT |
| 55 | SI: E55K (1) | TCT -> TTT |
| 60 | SI: D60N (7) | TCT -> TTT |
|  | SIV: D60H (7) | TCT -> TGT |
|  | SI: D60H (2) | TCT -> TGT |
|  | SIV: D60N (2) | TCT -> TTT |
| 61 | SI: E61Q (3) | TCA -> TGA |
|  | SI: E61K (1) | TCA -> TTA |
|  | SI: E61N (1) | TCA -> TTA |
| 62 | SIV: D62N (5) | TCA -> TTA |
|  | SIV: D62H (2) | TCA -> TGA |
| 69 | SIV: R69K (8) | TCT -> TTT |
| 73 | SI: E73Q (6) | TCA -> TGA |
|  | SIV: E73Q (4) | TCA -> TGA |
|  | SIV: E73K (2) | TCA -> TTA |
| 75 | SI: D75N (15) | TCA -> TTA |
|  | SI: D75H (1) | TCA -> TGA |
| 77 | SIV: E77Q (5) | TCA -> TGA |
|  | SIV: D77H (4) | TCA -> TGA |
|  | SII: D77H (3) | TCA -> TGA |
|  | SIV: D77K (1) | TCA -> TTA |
|  | SII: D77N (1) | TCA -> TTA |
| 82 | SI: E82Q (13) | TCT -> TGT |
|  | SI: E82K (3) | TCT -> TTT |
|  | SIV: D82N (3) | TCT -> TTT |
|  | SI: E82Q (1) | TCA -> TGA |
|  | SI: E82X (1) | TCT -> TGT |
|  | SI: E82Y (1) | TCT -> TGT |
|  | SI: D82Q (1) | TCT -> TGT |
| 83 | SI: R83K (17) | TCT -> TTT |
|  | SIV: R83K (1) | TCT -> TTT |
| 93 | SI: R93K (1) | TCT -> TTT |
| 101 | SII: E101K (1) | TCA -> TTA |
| 119 | SII: E119K (1) | TCT -> TTT |
| 138 | SIV: E138Q (5) | TCA -> TGA |
|  | SII: E138Q (1) | TCA -> TGA |
|  | SIV: E138H (1) | TCA -> TGA |
|  | SIV: E138H (1) | TCT -> TGT |
| 158 | SI: D158N (1) | TCT -> TTT |
| 175 | SI: E175Q (1) | TCT -> TGT |
|  | SIV: E175Q (1) | TCT -> TGT |
|  | SI: D175H (1) | TCT -> TGT |
| 215 | SI: R215K (1) | TCT -> TTT |
|  | SIV: R215K (1) | TCT -> TTT |
| 237 | SI: V237I (1) | TCA -> TTA |
| 258 | SI: D258N (1) | TCA -> TTA |
| 292 | SI: R292K (1) | TCT -> TTT |
| 340 | SI: R340T (1) | TCT -> TGT |

AA, amino acid; BCR, B cell receptor;
SI, serotype I; SII, serotype II; SIII, serotype III; SIV, serotype IV

The Vp1 pentamer homology model (**Figure 3**) allowed visualization of the *intra*-pentameric interactions depicted in green between chain 2 and 3 or the *inter*-pentameric interactions of chain-1’ with chain 4 and 5 (orange). Further, we depicted other relevant entities (**Figure 3**) such as sialic acid receptors (chain-2, blue), neutralizing antibodies (chain-3, pink), or the inner capsid protein Vp2/Vp3 (yellow). The *per* residue average QMEAN score of the combined Vp1 chain-1 to -5 together with chain 1’ is displayed in chain-1 (**Figure 3**). For the subsequent analyses, we focused on variant residues with a prevalence of more than 1% in GenBank or being present in our recent study (10) (summarized in **Table 3**).

**Figure 3.**
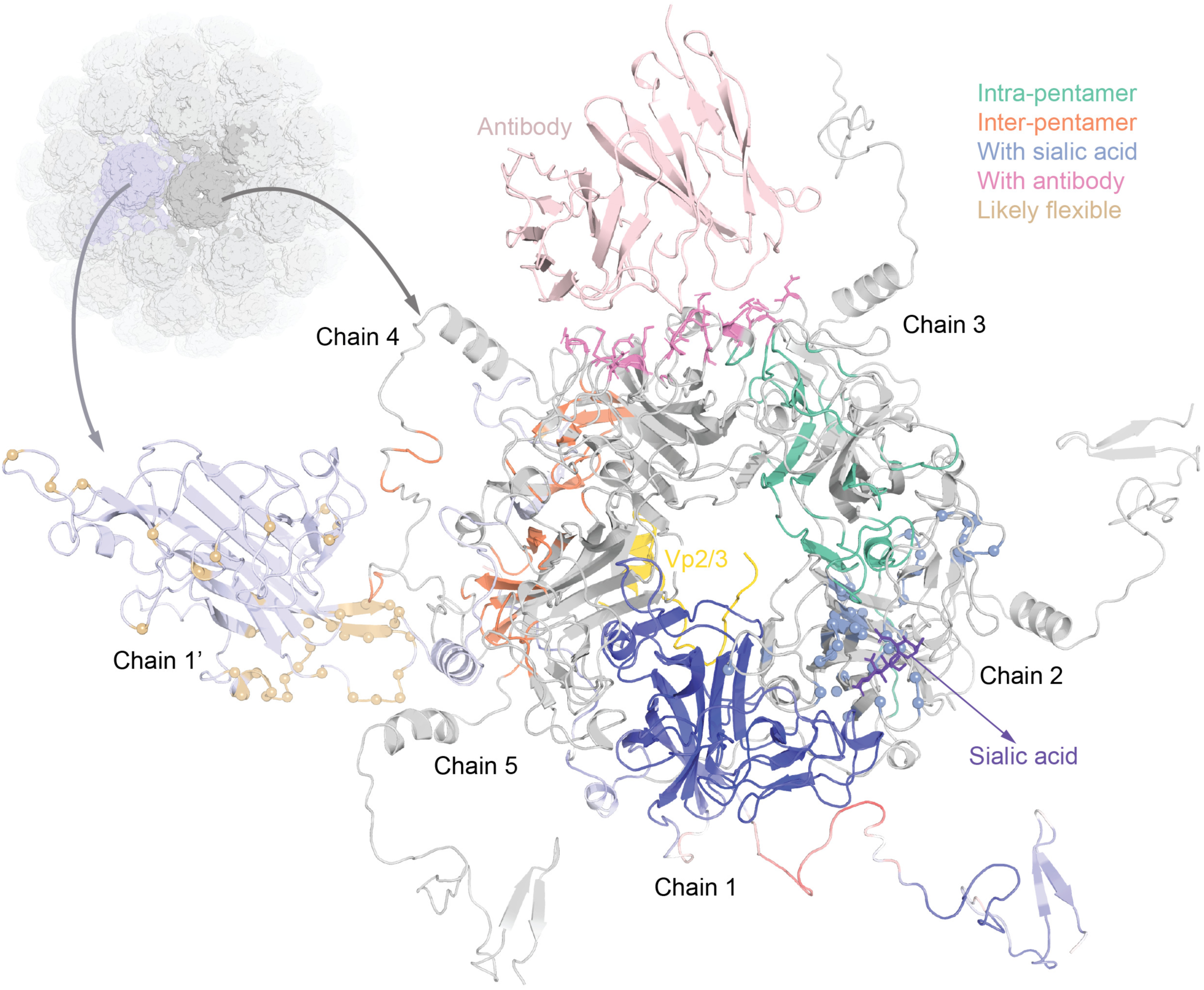
Vp1 protein structure and interaction predictions. All chains of one pentamer are shown and labelled counter-clockwise as chain-1 to -5, as well as the neighboring first chain of the second pentamer (labelled chain-1’), a sialic acid receptor (from PDB: 6ESB, in purple), a neutralizing antibody (from PDB: 7PA7, in light pink) and an interacting domain of Vp2 (from PDB: 1CN3, in yellow). Structure quality is depicted for chain-1 using QMEAN scoring. Contacts are colored on chains 2-5, residues with median EDIA_m_ <0.8 are shown as spheres and colored on chain-1’.

**Table 3.**
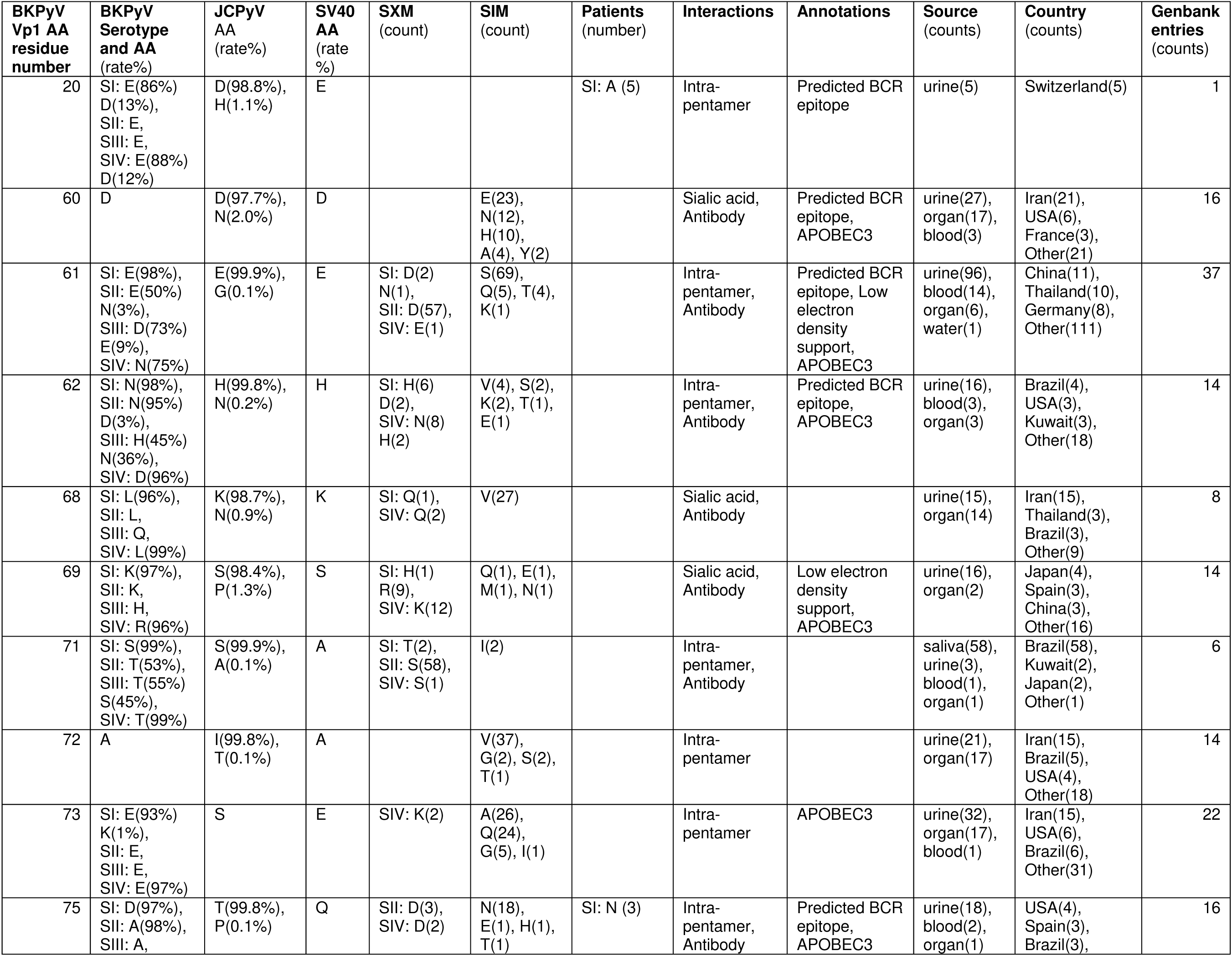
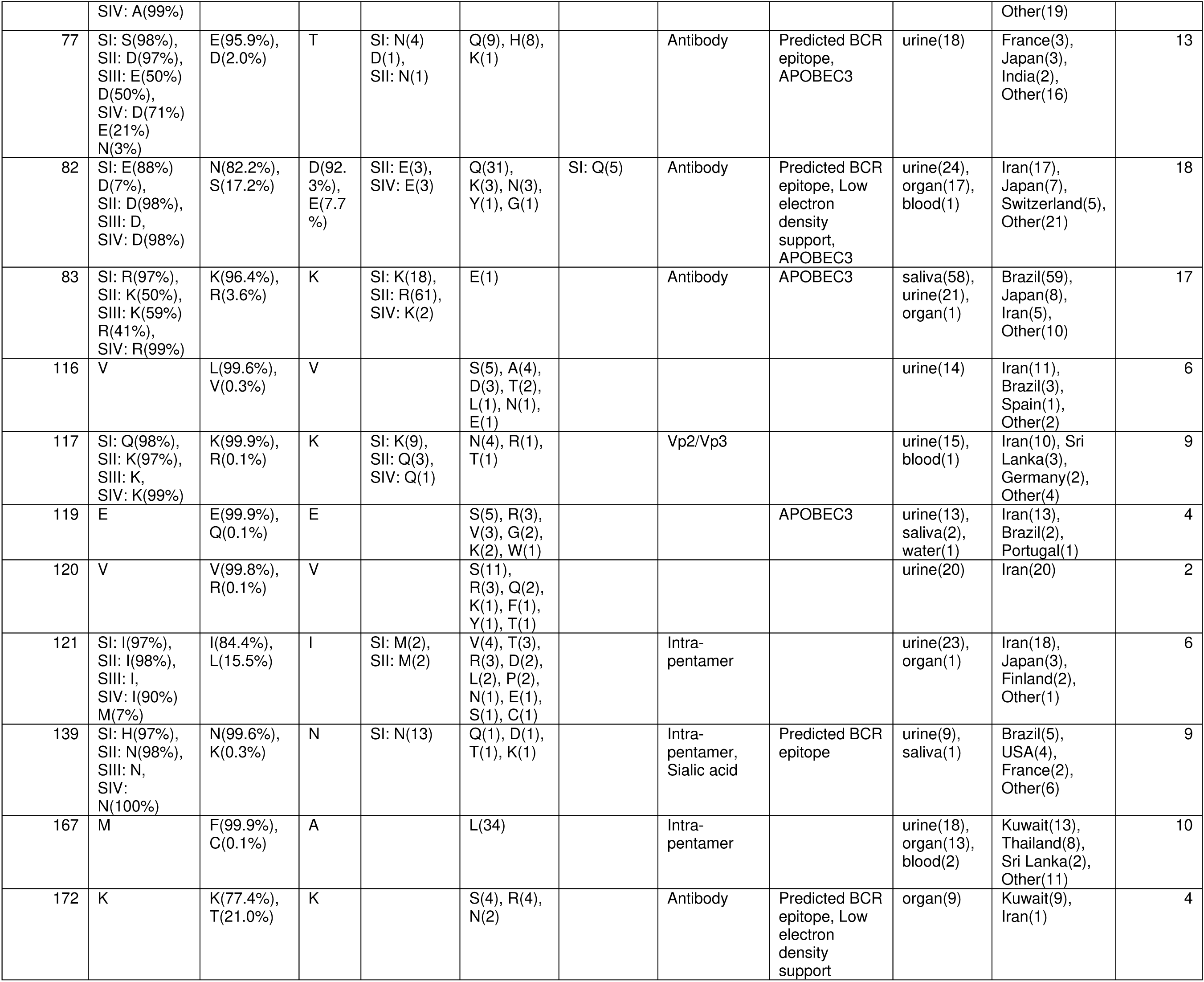
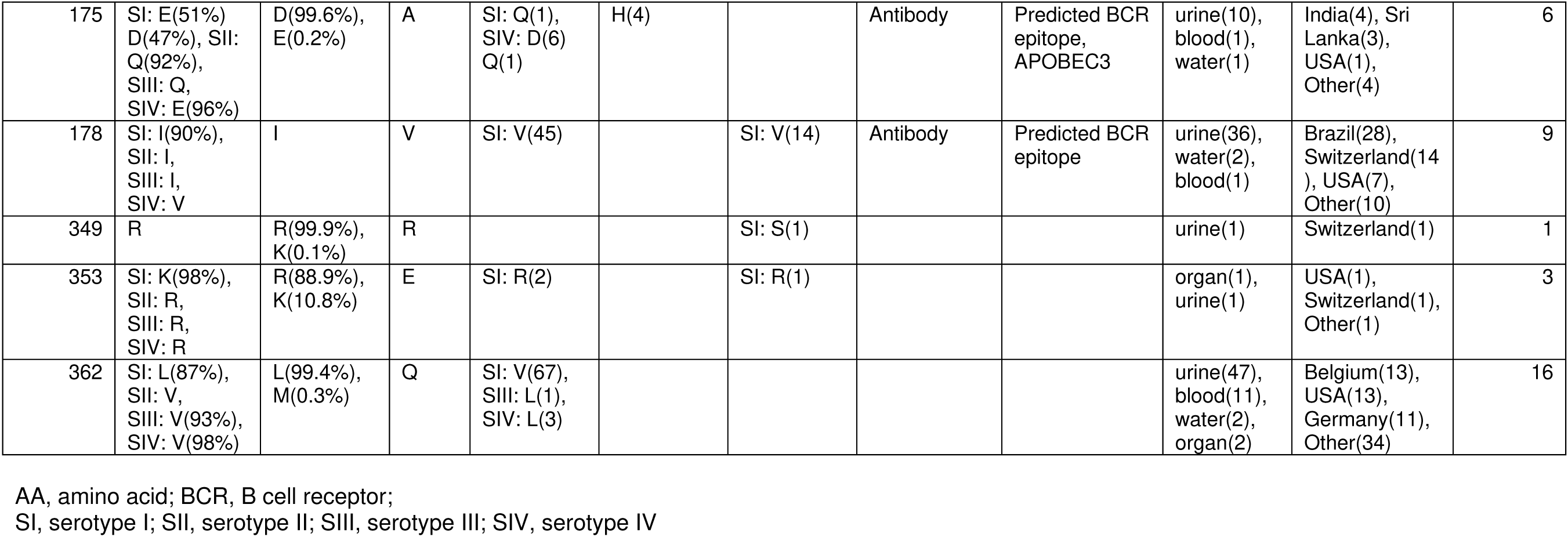
Summary of notable Vp1 variants.

A comparison of 9 available structures (see Materials & Methods) of the BKPyV-Vp1 revealed highly similar conformations of the BC-, DE-, and HI-loop, with a median Cα distance of <1Å across all residues (**Supplementary Figure 1**). Parts of Vp1 at the N-terminus and C-terminus were found to be less structurally conserved (>2Å distance), and six of nine structures only covered the Vp1 core and not the N- or C-terminal regions. As the available structures of the BC-, DE-, and HI-loop might not capture their inherent flexibility, we used the underlying electron density data of 3 BKPyV crystal structures (PDB IDs: 4MJ0, 4MJ1 and 7ZIQ) to pinpoint amino acid residues with low average electron density support, defined by a median electron density score (EDIA_m_) < 0.8 (29) (**Figure 2, bottom panel**, light brown). These more flexible residues clustered in three regions, around amino acid 40, 100 and 300 of the BKPyV-Vp1 protein shown in the structural model (**Figure 3, depicted on chain-1’,** light brown), in agreement with the regions of >2Å across available structures (**Supplementary Figure 1**). However, we noted four residues at positions 61, 69, 82, and 84 in the BC-loop and three residues at position 135, 138, and 143 in the DE-loop also had low density support pointing to higher flexibility (**Figure 2**). Indeed, SXM and SIM were mostly found at these four residues in the BC-loop (**Figure 2, center panel**).

#### Distribution and variation of Vp1 SDPs and SXMs

The frequency of serotype-defining amino acids found in the respective SDPs among the available sequences are shown on the 3D model of a Vp1 monomer (**Figure 4, spheres,** position, and serotype-I, -II, -III, and -IV from top to bottom). The SDPs were spread across the Vp1 structure but were enriched in the BC-loop (12/29; dark brown spheres) in line with its role as the major serotype-defining domain. For 18 SDPs, three out of four serotypes appeared to be conserved, with only one serotype having a different amino acid, usually serotype I (11/18) or serotype IV (4/18). In two-thirds of SDPs, the serotype-defining amino acids had similar physicochemical properties for the four serotypes (**Figure 4**), being positively charged (residues 69, 83, 215, 316, 353), negatively charged (residues 20, 82, 138), polar (residues 71, 74), non-polar (residues 42, 145, 178, 210, 362), or bulky (residue 66). Most SDPs were found in loops, in line with their being able to accommodate some variation. Conversely, only three SDPs (residues 117, 210, and 225) were located in structured beta-sheets.

**Figure 4.**
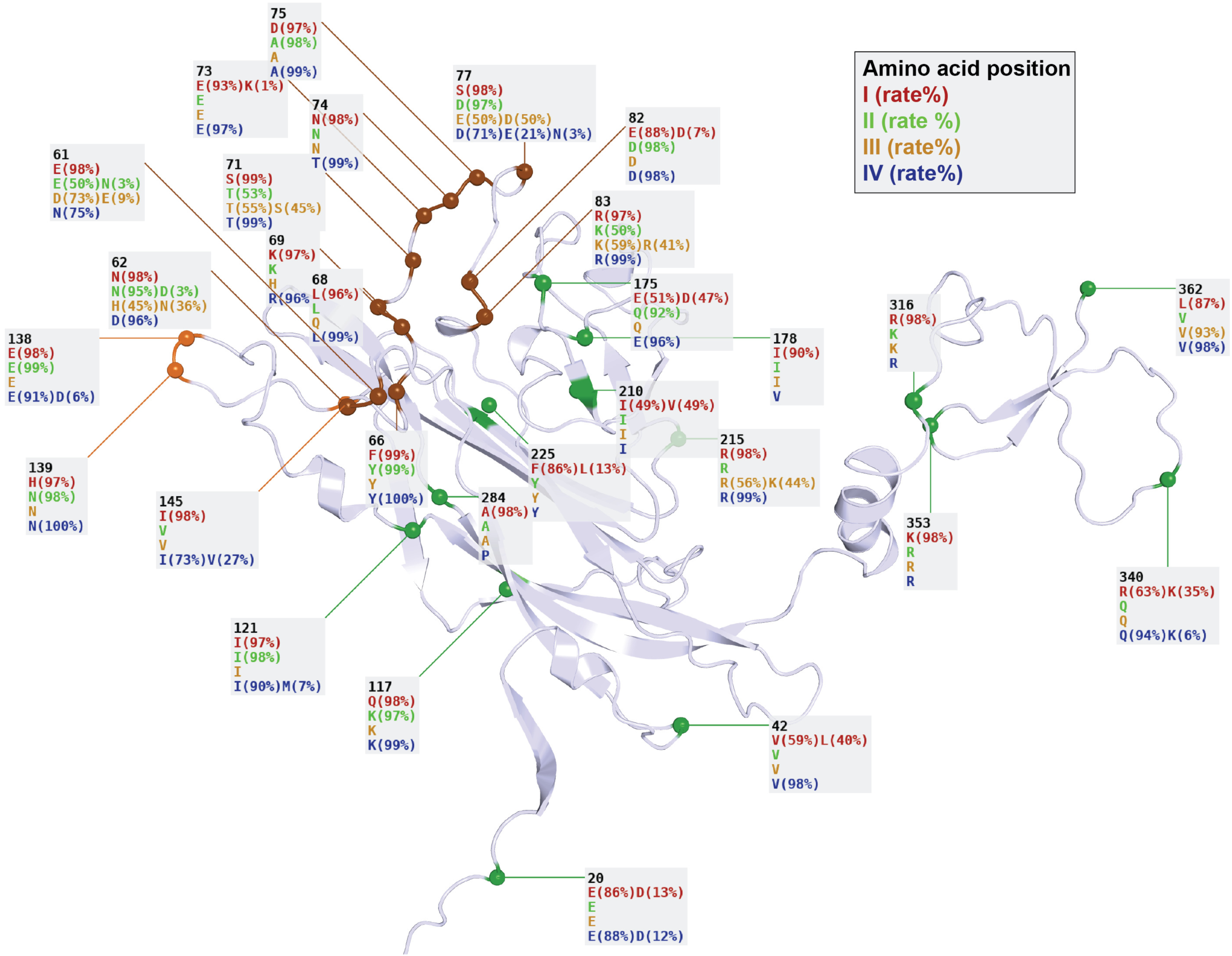
Serotype-defining positions depicted on the Vp1 structure. Each SDP is labelled with the residue position followed by the amino acids found in the serogroup reference sequences of serotypes I, II, III and IV respectively. The percentage of each amino acid across Vp1 GenBank entries assigned to that serotype is shown (blank = 100%). Residues in the BC and DE loops are colored.

To investigate the potential impact of the variant amino acids in the Vp1 protein, we depicted the SXM and SIM present in at least three entries in the context of the 3D structure of the Vp1 monomer (**Figure 5**). The most prevalent SXMs in serotype-I were L362V (67 entries) and I178V (59 entries) reflecting 6.2% and 5% of all serotype-I assigned entries, respectively. In both cases, the SXM represented conservative non-polar exchanges. For serotype-IV, the most prevalent SXM were R69K (12 entries, 4.1%) and D62N (8 entries; 2.74%). R69K represented a conservative positively charged exchange. The 3D structural information suggested interaction of residue 69 with sialic acid receptors or neutralizing antibodies, while the low electron density support indicated that the size difference of R69K is likely to be well accommodated. In contrast, D62N represented a non-conservative change in charge and size which is notable because of its prominent position regarding intra-pentameric and neutralizing antibody interactions. Both of these SXM are also possible APOBEC3 mutations (**Table 2**). For serotype-II, the most prevalent SXM were T71S (58 entries) and K83R (61 entries), both representing conservative changes at the C-terminal end of the BC-loop whereby both contacted neutralizing antibodies and the former also participated in intra-pentameric interactions. Notably, this high number of entries with both SXM were derived from a single study on saliva of human subjects from Brazil, which made up 47% of the rather uncommon serotype II. When including entries from other parts of the world, the SXM E/N61D was notable with 57 entries, where the same interaction considerations apply as for residue 62 (**Figure 5**). For serotype III, no SXMs were found above our chosen frequency threshold of 1% which might reflect the scarcity of serotype III-assigned entries. However, when counting serotype II-SXMs, three Vp1 positions distinguishing serotype II and III (SDPs 61, 71, and 83) had the same amino acids, indicating that these two serotypes differ only in positions 62 and 77.

**Figure 5.**
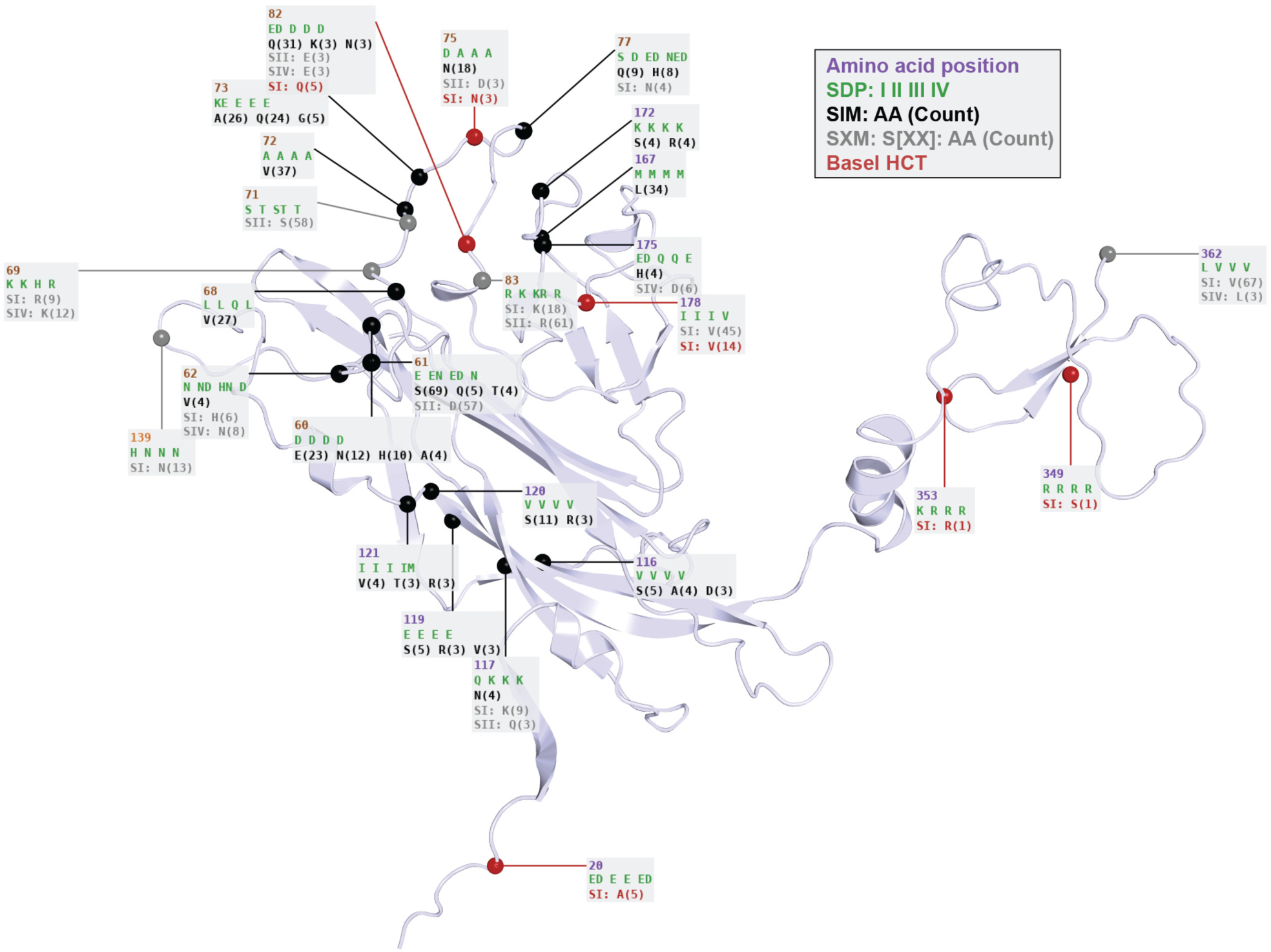
Serotype-exchange and serotype-independent mutations occurring with over 1% frequency or in the Basel HCT study depicted on the Vp1 structure. Each mutation is labelled with the residue position, the amino acids found in serotypes I, II, III and IV at that position (in green), GenBank serotype-independent amino acids and counts for counts over 3 (SIM, in black), GenBank serotype-exchange counts for each serotype for counts over 3 (SXM, in grey, with serotype labelled), and changes in Basel HCT patient samples (in red, labelled with the serotype they are found in for both SXM and SIM).

#### Structural impact of sequence variation in the Vp1 BC-loop

The combined data indicated that the BC-loop region (residues 59-83) contained the most relevant information for serotyping as shown by a high number of SDPs, high rates of SXMs but also SIMs (**Figure 5**). Accordingly, the serotype-defining amino acids within this loop had differing physicochemical properties, already indicating its potential for non-conservative SIMs. A number of SIMs were found at residues 72 (A) and 73 (E/K), with 57 entries featuring one of the following combinations: V_72_A_73_, A_72_Q_73_, V_72_Q_73_, A_72_A_73_, V_72_G_73_, and V_72_E_73_. All these combinations allowed for the BC-loop to adopt a different conformation compared to the reference. Indeed, previous reports suggested that mutations at positions 72 and 73 could lead to BC-loop flipping (45), specifically with E73A alone and E73Q combined with A72V.

#### Structural impact of Vp1 variants on sialic acid binding

A set of well-documented variant residues at positions 69, 72, 73, and 82 in the BC-loop, and 139 in the DE-loop, have been recognized for their role in sialic acid binding (**Figure 6A**). These also represent some of the most prevalent SIMs in our analysis. Residues 69 and 82 both display low electron density support in crystal structures, suggesting high flexibility in that region. In general, the extensive variation observed in these sialic acid binding residues may contribute to differences in glycan specificity. Indeed, a mutation in residue 69, also in combination with residue 82, has been shown to alter or eliminate glycan specificity (45, 46). Thus, mutations in these residues would be impacting host cell specificity and infection.

**Figure 6.**
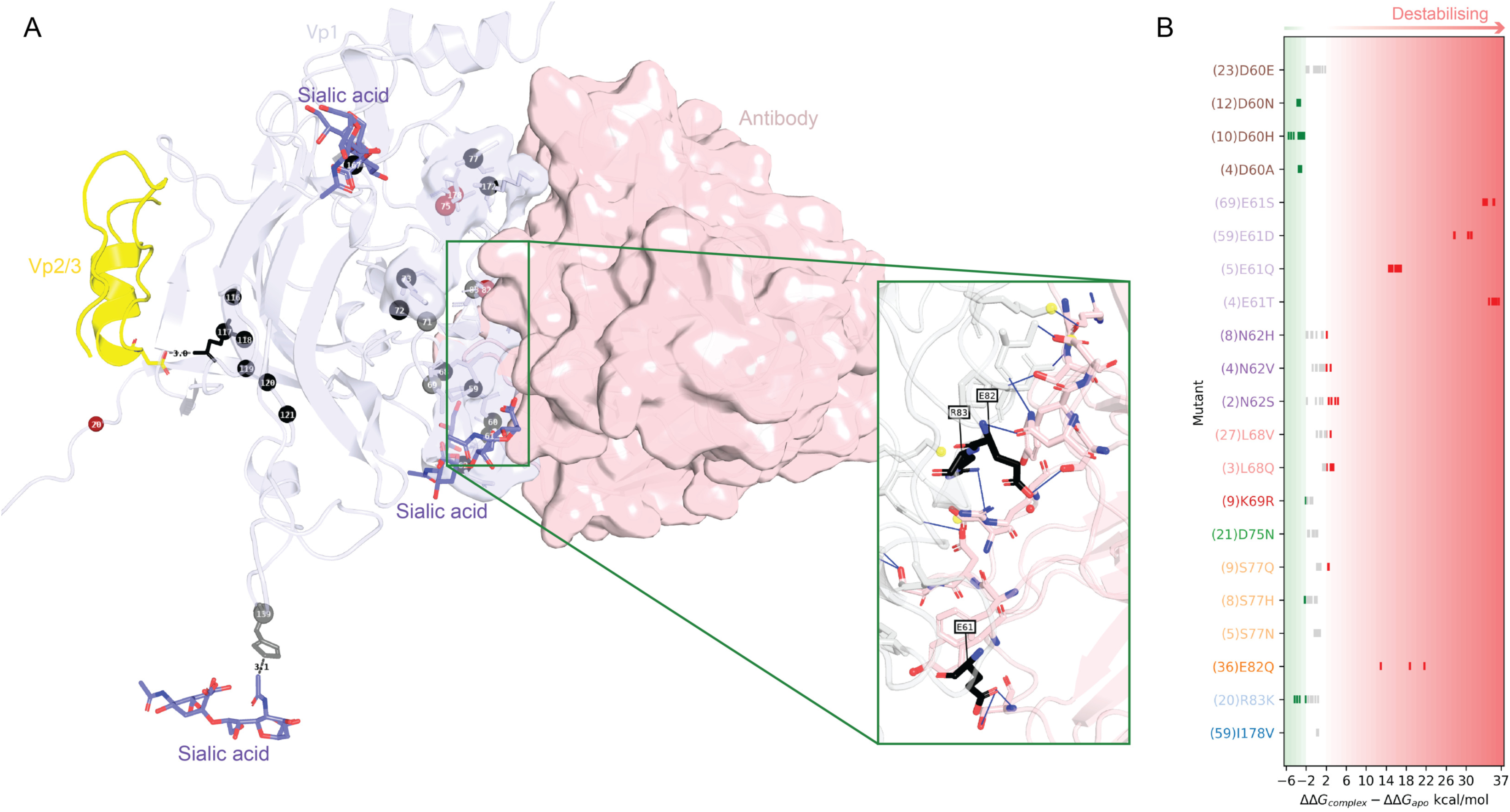
Vp1 interactions with sialic acid receptors, Vp2/Vp3 and neutralizing antibodies. A. Vp1 monomer (in grey) in in complex with scFv 29B1 antibody (in light pink, from PDB: 7PA7). Variant residues are labelled and hydrogen bonds between Vp1 residues and antibody residues are shown as blue lines in the inset. Sialic acid receptors (from PDB: 6ESB) and the Vp2/Vp3 fragment (from PDB: 1CN3) are shown in purple and yellow respectively. B. FoldX mutagenesis analysis of commonly found antibody-binding mutants. The difference in predicted ΔΔG values of the mutated complex (PDB: 7PA7) with and without the antibody is shown for 3 replicates of each (9 differences), with red representing predicted destabilization and green predicted stabilization of the interaction.

However, since infection can be maintained in principle despite altered, reduced, or lost glycan binding (45), mutations in this region might have a limited evolutionary cost. This hypothesis could also apply to residues 60 and 68 both of which had numerous SIMs leading to significant alterations in amino acid properties and thus sialic acid binding (**Figure 5 and 6A**).

#### Impact of Vp1 variants on protein interactions

The region preceding the DE-loop (residues 116-121) exhibited considerable variation within BKPyV-Vp1 but appeared to be more conserved in the corresponding Vp1 positions of JCPyV and SV40 (**Figure 2**, top panel). However, this variation is mostly derived from four studies examining *VP1* sequences in Iranian patients, while these residues were infrequently affected in sequence entries from other countries (**Table 3**). A noteworthy variant is residue 117, which in other polyomaviruses is a lysine and believed to form a salt bridge/hydrogen bond network with an aspartate in the minor capsid proteins Vp2 or Vp3 (**Figure 6A**). NMR studies have revealed that this residue also interacts with a Vp2/Vp3-like peptide in BKPyV (47). In BKPyV serotype I, this residue is a glutamine, but the identified variants included lysine, histidine, and arginine, all of which are able to form salt bridge/hydrogen bonds.

In the Basel HCT study, five samples had an alanine at residue 20 at the otherwise highly conserved aspartate or glutamate at the base of the pentamer, likely playing a role in capsid formation. This is especially interesting, since, in general, residues involved in inter-pentameric interactions (**Figure 2, orange**) have much lower mutation rates (0.12 ± 0.12 %) compared to the rest of the Vp1 sequence (0.3 ± 1 %). We concluded that changes affecting protein-protein interactions can occur in principle in the BKPyV capsid but seem to be rare.

#### Vp1 immunogenic epitopes and potential for immune escape

To investigate changes of potential relevance for adaptive immune control, we employed *in silico* epitope prediction approaches to predict B-cell receptor (BCR) epitopes from the Vp1 structural model and HLA-class I epitopes for the most common HLA-A, -B, and -C types from the Vp1 linear sequence (**Figure 2**). The predicted HLA-I epitopes were spread over the entire Vp1 sequence. However, the epitopes predicted across the widest range of HLA types were often near the end of the Vp1 sequence (starting at positions 302, 332 and 338) where SXMs and SIMs are rare (**Figure 2, center and bottom panels**). One epitope within the highly variable BC-loop region, S_78_SDSPERKM_86_ was predicted to bind to multiple common HLA-C types, while the remaining epitopes were rather specific for the specific alleles. Overall, epitopes were predicted for at least one common HLA type for all the mutation hot spots found, indicating the potential of some immune escape according to host HLA type.

The BCR epitope prediction agreed with previously determined antibody-binding regions (from PDB IDs 6GG0 and 7PA7), but also indicated potentially novel BCR epitopes which have not been structurally characterized so far, namely in the DE-loop (residues 135-138) and HI-loop (residues 274, 275) (**Figure 2**). Half of the prevalent variant residues (13/26) were located in NAbs-binding regions (**Figure 6A, gray surface**). Using FoldX (36), we compared the predicted energies upon mutating these residues in antibody-free and antibody-bound structures (**Figure 6B**). This revealed that the observed variants in residues 61 and 82, residues forming hydrogen bonds with the antibody (**Figure 6A** inset), are highly likely to destabilize antibody binding and thus could lead to immune escape. Mutations allowing for evasion of neutralizing antibodies and T cell response are especially likely to confer a selective advantage. Thus, a number of the prevalent variant positions seen in our analysis are likely to be associated with immune evasion which can evolve and hence delay BKPyV-specific immune reconstitution in immunosuppressed patients.

### Diversity of the BKPyV regulatory LTag

The 695 amino acid-long LTag protein is commonly divided into five domains (**Figure 7A rectangles**), for which we predicted the 3D structure to visualize the location of variant amino acids (**Figure 7B-D**). The domains are the DnaJ (**Figure 7A and 7B blue**), the retinoblastoma protein binding (pRb) (**Figure 7A, orange,** unstructured), the origin binding domain (OBD) for viral genome replication, **Figure 7A and 7C, light green**), the helicase (**Figure 7A, 7C and 7D, purple and gray)**, and the host range (HR) domain (**Figure 7A peach,** unstructured). The N-terminal DnaJ domain is identical for LTag and small T-antigen, which are derived by splicing. The helicase domain is further divided into three structural domains (D1-D3), of which D3 is discontinuous, consisting of D3-1 and D3-2. Residues participating in intra-helicase interactions are depicted (**Figure 7A, purple triangles**). The helicase domain binds to DNA in conjunction with the OBD (**Figure 7C**), and then forms a hexamer (**Figure 7D**). We depicted residues participating in the interaction of one LTag monomer with DNA, ATP, zinc ions (**Figure 7C**) as well as the interaction of one monomer of the LTag hexamer to the tumor suppressor p53 protein, which may contribute to oncogenic transformation (**Figure 7D**).

**Figure 7.**
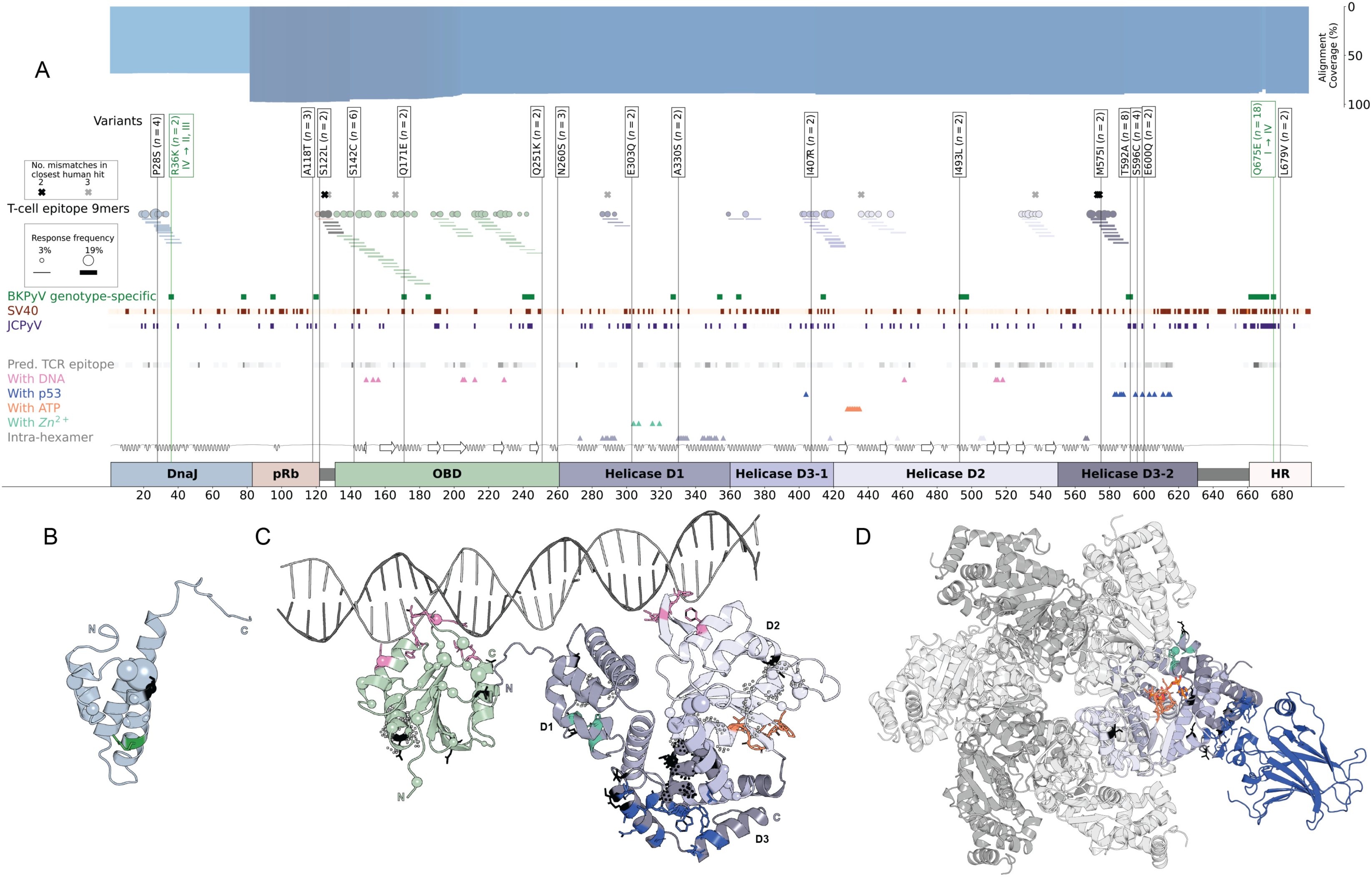
Variability and binding in BKPyV large tumor antigen protein. A. BKPyV LTag domains are shown across the entire sequence length, with secondary structure depicted above (loop = line, sheet = arrow, helix = wave). The LTag Helicase is further divided into D1, D2 and D3 domains. Colored triangles mark residues involved in helicase intra-hexamer interactions and those binding to Zn^+2^, ATP, DNA and p53 as seen in SV40 LTag. Dark green squares mark genotype-defining residues. Purple and orange lines mark residues where none of the amino acids in BKPyV reference sequences are found in corresponding positions in the JCPyV and SV40 LTag entries, respectively. HLA-presenting T-cell 9mer epitopes are labelled as lines along the sequence, with their starting positions as circles along the sequence. Line thickness and circle size correspond to the 9mer epitope response frequency in kidney transplant patients (12, 23, 24). Black and gray crosses mark the starting positions of 9mer epitopes which have a match in the human proteome with 2 and 3 mutations respectively. Genotype-exchange and genotype-independent mutations found in BKPyV LTag GenBank entries are labelled with the count and amino acid change and colored black in the structure. B. AlphaFold structural model of LTag DnaJ domain C. AlphaFold structural models of LTag OBD and Helicase domains in complex with DNA (from PDB ID: 4GDF) D. AlphaFold model of LTag Helicase arranged in a hexamer (as in PDB ID: 1SVM), depicting a Zn^+2^ residue in green (from PDB ID: 1SVM), ATP molecule in orange (from PDB ID: 1SVM) and p53 protein in blue (from PDB ID: 2H1L). In panels B, C, and D, interacting residues are shown as colored sticks following the legend in panel A, GenBank variations as black (for genotype-independent) and green (for genotype-exchange) sticks, the starting residues of confirmed 9mer epitopes as spheres with the radius corresponding to KT patient frequency, and peptide regions with matches in the human proteome as black (for 2 mutations) and gray (for 3 mutations) dots.

We identified and analyzed 742 BKPyV-*LTAG* entries in GenBank which corresponded to 169 unique protein sequences. Coverage of the LTag protein by these sequences over 90%, except for 70% the DnaJ domain (**Figure 7A, top panel,** blue bar). Overall, our analysis revealed a higher degree of conservation compared to the *VP1* entries, with only 17% exhibiting variations from the assigned genotype, and only 16 amino acid variants supported by at least two *LTAG* entries (**Figure 7A, top panel,** boxes). Three of these 16 changes (P28S, S122L, and I407R), were non-conservative. BKPyV LTag sequences differing from both JCPyV and SV40 LTag were mainly located in the helicase and HR domain (**Figure 7A top panel,** red and purple squares). Conversely, the overview plot identifies LTag regions which are conserved between BKPyV, JCPyV and SV40, and might be potential candidates for cross-protective immunogenic epitopes.

Next, we mapped location and response rate for 97 BKPyV immunodominant CD8 T cell 9mer epitopes previously identified in kidney transplant patients (12, 23, 24) (**Figure 7A, top panel,** circles and lines show the 78 confirmed epitopes). To investigate whether or not the immunodominant 9mer epitopes had any similarities with proteins of the human proteome, we conducted a peptide sequence search with PepMatch (see Materials & Methods). While none were identical, three peptides had two mismatches and six peptides had three mismatches compared to the human counterpart (**Figure 7**, black and gray crosses respectively for two and three mismatches). Specifically, the viral 9mer S_125_**T**PPKKKR**K** matched the S**K**PPKKKR**R** of the “G patch domain-containing protein 4” sequence with non-conservative amino acid change at the anchor position 2 and a conservative change at the anchor position 9. The viral 9mers **S**_573_G**M**TLLLLL and G_574_**M**TLLLLL**I** matched the **P**G**W**TLLLLL and G**W**TLLLLL**L** of “Leucine-rich co-lipase-like protein 1” sequences with nonconservative changes at anchor position 2. Notably, the viral 9mer S_573_**G**M**T**LLLLL matched the “Macrophage colony-stimulating factor 1 receptor” sequence S**I**M**A**LLLLL with a change at anchor position 2.

The most frequent 9mer epitope in kidney transplant patients, L_27_PLMRKAYL_35_, falls within the DnaJ region of LTag which is also shared with sTag. However, it is found mutated in 4 BKPyV entries (P28S), which abolishes predicted binding to 36/44 common HLA-B51, B7, B8, and A24 alleles (12, 48). It also has variations in JCPyV and SV40 sequences (L27I in 196 JCPyV and 22 SV40 entries, and L29V in 186 JCPyV entries), though the predicted HLA binding is not affected by these changes. In contrast, epitopes K_216_LCTFSFLI_225_ and C_218_TFSFLICK_227_ with response frequency of 8.0% and 5.5% in kidney transplant patients, respectively, are conserved in all available sequence entries for BKPyV, JCPyV and SV40. K_216_LCTFSFLI_225_ is predicted to bind to 33 HLA-A2, A32 and B13 alleles and was found in patients having HLA-A2 and HLA-A24. C_218_TFSFLICK_227_ is predicted to bind to 41 HLA-A11, A30, A31, A68, A03, A34, A66 and A74 alleles and was found in patients having HLA-A3. Of note, we could not identify matching epitope hits in the human proteome.

## DISCUSSION

Type and rate of amino acid variations in BKPyV may provide important insights into BKPyV diversity in human populations and a first step towards better defining determinants of BKPyV-specific immunity needed to protect vulnerable patients from BKPyV diseases. Indeed, our earlier work identified BKPyV mutant epitopes downregulating polyfunctional cytotoxic T cell activation (10, 11). Similar changes in serotypes could predict escape from neutralizing antibody responses (49, 50). Here, we present a comprehensive analysis of amino acid variations in publicly deposited sequences of the BKPyV Vp1 and LTag proteins and include a cross-check with recent data from our center. Using available experimental structures and computational modelling, we placed the amino acid variations in their structural context to visualize conformational, functional, or immunologic aspects.

Our study provides the following insights: first, BKPyV-genotype(gt)1 was found in 71.2% of publicly deposited Vp1 GenBank entries, followed by BKPyV-gt4 (19.3%), BKPyV-gt2 (8.1%) and BKPyV-gt3 (1.4%), but prevalence rates differed according to geography and specimen type. Second, 43% of Vp1 carried serotype-exchange mutations (SXMs) or serotype-independent mutations (SIMs) whereby 18% had more than one amino acid mutation and included changes in antibody-binding domains. Third, LTag sequences were largely conserved, with only 16 mutations detectable in more than one entry and typically without significant effects on LTag-structure or interaction domains. However, some LTag changes were predicted to affect HLA-class I presentation of immunodominant 9mers to cytotoxic T-cells.

Overall, Vp1 sequences displayed a high degree of amino acid variability, suggesting a remarkable plasticity of the major capsid protein, with mutation hotspots around the exposed BC-and DE-loops, which include SDM and neutralizing domains and clearly differ from JCPyV and SV40. There were numerous SXM in positions defining the four major Vp1 serotypes, suggesting that the existing set of serogroup reference sequences may not adequately cover the common serotype-specific contexts. Of note, sequences assigned to different serotypes differed in SXM rates, whereby 95% of serotype-II assigned sequences had at least one SDP while this percentage was lower with 17%, 4.5%, and 14% for serotypes-I, -III, and -IV, respectively. One SXM has been shown to be a neutral mutation within a specific serotype context (N61S within serotype IV) (16). The relative structural flexibility of common SXM positions implies the potential of accommodating additional neutral mutations. Previous studies have suggested combining BKPyV-serotype II and III into a single serotype (51). Our own observations demonstrate that BKPyV-serotype II and III sequences differ in only two Vp1 positions (62 and 77). Combining both serotypes into a new II-III would be strengthened by further support through independent sequence entries. More importantly, functional studies are needed characterizing serological or functional cross-reactivity as well as shared properties regarding viral infection, replication and neutralization to justify a novel combined serotype II-III classification. Further, a majority of 834/1286 (65%) Vp1 variants were SIM and hence not linked to a particular serotype. However, the rates of SIMs differed in the different serotypes, being highest with 38% for serotype IV compared to 25%, 20%, 27% for serotypes I, II, and III, respectively. The sequences from our HCT study were mainly from serotype I (63/65 sequences) and contained common SIM (D75N, E/D82Q) and SXM (I178V), except for E/D20A which was not observed in the public data.

Of 26 prevalent variants found in Vp1, our structural studies revealed that 6 concerned interactions with sialic acid receptors, 13 changes concerned sites contacting antibodies, and 10 were potentially involved in intra-pentamer interactions. In contrast, Vp2/Vp3 interaction was only indicated in 1 case. These data suggest that the outer surface of the BKPyV virion can accommodate more variability compared to the inner surface and may not only contribute to serotype specificity but also to gaps in the humoral defense. Indeed, mutations in residues 61 and 82 are predicted to destabilize antibody binding, and those in residues 60, 58, 69, 72, 73, and 82 can alter glycan binding. Residue positions 61, 69, 82 and 172 had low electron density support in experimentally solved crystal structures indicating their potential for flexibility. Interestingly, for eight common variant positions, more than one sequence displayed nucleotide changes matching the APOBEC3 mutational signature. This includes the SIMs D60H/N, E61Q, E73Q, D75N, D77H, E77Q, E82Q/K and D82N; serotype-IV SXM D62H/N, R69K; and serotype-I SXM R83K. Some of these (D62N, R69K, and D77H) have been identified as APOBEC3 mutations in previous studies (15). APOBEC3 belongs to a family of ssDNA cytosine deaminases that has been linked to innate antiviral defense by mutating viral genomes (52). APOBEC3 mutations have been observed across a range of viruses, though the level of mutation appears to be significantly lower in DNA viruses compared to retroviruses (53). However, APOBEC3-like signature mutations were not found at all in 15 prevalent variant BKPyV positions. Taken together, based on structural context, we identified a number of variants with different interactions, which could have significant implications for BKPyV replication, virion interaction with host cells and the adaptive immune response.

In contrast, our LTag sequence analysis revealed higher conservation in line with its central multi-functional role often compared to a Swiss army knife, coordinating polyomavirus replication together with timed recruitment of essential host cell functions. Of the only 16 amino acid variants reported at least twice in 695 amino acids, each of the relevant domains had at least one change, whereby possibly the helicase, and in particular the helicase D3-2 appeared to be slightly more affected. Our structural analysis suggested that the amino acid variants were not associated with major conformational changes in line with its conserved multifunctional role. Nevertheless, several variants were non-conservative with respect to size or charge. While these variants are less dramatic than the ones known for the chromosomally integrated Merkel cell PyV in cases of Merkel cell carcinoma, which include protein truncations and/or frame shifts, the functional consequences of LTag mutations on BKPyV replication, if any, need to be addressed in relevant infection models.

Nevertheless, our analysis revealed potential immunological consequences of the LTag amino acid variants. Indeed, the variant changes affected the type and binding to HLA-class I of immunodominant CD8 T cell 9mer epitopes previously identified in kidney transplant patients (12, 23, 24). In particular, the 9mer epitope L_27_PLMRKAYL_35_ in the DnaJ region of LTag was mutated in 4 BKPyV entries (P28S), which abolishes predicted binding to 36/44 common HLA-B51, B7, B8, and A24 alleles (11, 12, 48) and which was reported to contribute to protection of kidney transplant recipients from BKPyV-DNAemia (48, 54). The BKPyV 9mer epitope L_27_PLMRKAYL_35_ homologue also has variations in JCPyV and SV40 sequences (L27I in 196 JCPyV and 22 SV40 entries, and L29V in 186 JCPyV entries), though the predicted HLA binding is not affected by these changes. In contrast, epitopes K_216_LCTFSFLI_225_ and C_218_TFSFLICK_227_ with having rates of 8% and 5.5% in kidney transplant patients, respectively, are conserved in all available sequence entries for BKPyV, JCPyV and SV40. K_216_LCTFSFLI_225_ is predicted to bind to 33 HLA-A2, A32 and B13 alleles and was found in patients having HLA-A2 and HLA-A24. C_218_TFSFLICK_227_ is predicted to bind to 41 HLA-A11, A30, A31, A68, A03, A34, A66 and A74 alleles and was found in patients having HLA-A3.

Comparison of LTag with the human proteome revealed only a few matches and were often of low complexity with repetitive amino acids, such as leucine in S_573_GMTLLLLL and lysine in S_125_TPPKKKRK. Conversely, we identified two immunodominant epitopes conserved across BKPyV, JCPyV and SV40, which may be highly relevant targets for immunotherapy and vaccine design. These epitopes lie within the DNA-binding region of the LTag OBD required for the viral genome replication, the mutation of which may significantly increase the evolutionary cost of immune escape.

Limitations of our study are the reliance of the analysis on publicly deposited Vp1 and LTag sequences for BKPyV, SV40 and JCPyV. While differences in the portions of Vp1 and LTag sequenced by different studies resulted in uneven alignment coverage across the sequence, at least 50% of the analyzed sequences cover the entire protein, suggesting that the hotspot variability plots still provided a relevant representation of the possible variation. However, there were a few variants and mutant profiles which largely arose from sequences derived from a single study in a single country rather than multiple geographic locations, studies as well as healthy donors and affected and non-affected patients. Nevertheless, when including data from our own next generation sequencing study of HCT recipients in Basel, independent confirmation as well as new variants could be identified. Thus, these global data strengthen our earlier single-center findings detecting and functionally analyzing mutant 9mer epitopes meidating immune escape from HLA-I cytotoxic T cells. Taken together, this perspective and our analysis of the currently available BKPyV sequences reveal an unexpectedly high genetic variability for this double-stranded DNA virus that strongly relies on the host cell DNA replication machinery with its potential access to proof reading and error correction mechanisms. This should be taken into account when designing further approaches to antivirals and vaccines for patients at risk of high-level BKPyV replication due to insufficient virus-specific immunity.

---

**Supplementary Figure 1. Vp1 conformational differences across available structures** Superposition of all 58 chains of 9 Vp1 structures (PDB IDs: 7ZIQ, 6GG0, 6ESB, 7B6C, 7B6A, 4MJ1, 5FUA, 7B69, 4MJ0), colored by median Euclidean distance of corresponding Cα positions at each residue. The BC, HI and DE loops are indicated with dashed lines.

## REFERENCES

1. Torres C. 2020. Evolution and molecular epidemiology of polyomaviruses. Infect Genet Evol 79:104150.

2. Leuzinger K, Hirsch HH. 2023. Human polyomaviruses, p 2093-2130. *In* Carroll KC, Pfaller MA, Karlowsky JA, Landry ML, McAdam AJ, Patel R, Pritt BS (ed), Manual of Clinical Microbiology, 13th ed. ASM Press, Washington, DC.

3. Knowles WA, Pipkin P, Andrews N, Vyse A, Minor P, Brown DW, Miller E. 2003. Population-based study of antibody to the human polyomaviruses BKV and JCV and the simian polyomavirus SV40. J Med Virol 71:115–123.

4. Egli A, Infanti L, Dumoulin A, Buser A, Samaridis J, Stebler C, Gosert R, Hirsch HH. 2009. Prevalence of Polyomavirus BK and JC Infection and Replication in 400 Healthy Blood Donors. J Infect Dis 199:837–846.

5. Kean JM, Rao S, Wang M, Garcea RL. 2009. Seroepidemiology of human polyomaviruses. PLoS Pathog 5:e1000363.

6. Cesaro S, Dalianis T, Hanssen Rinaldo C, Koskenvuo M, Pegoraro A, Einsele H, Cordonnier C, Hirsch HH, ECIL-6 Group. 2018. ECIL guidelines for the prevention, diagnosis and treatment of BK polyomavirus-associated haemorrhagic cystitis in haematopoietic stem cell transplant recipients. J Antimicrob Chemother 73:12–21.

7. Imlay H, Xie H, Leisenring WM, Duke ER, Kimball LE, Huang ML, Pergam SA, Hill JA, Jerome KR, Milano F, Nichols WG, Pang PS, Hirsch HH, Limaye AP, Boeckh M. 2020. Presentation of BK polyomavirus-associated hemorrhagic cystitis after allogeneic hematopoietic cell transplantation. Blood Adv 4:617–628.

8. DeCaprio JA, Imperiale MJ, Hirsch HH. 2021. Polyomaviridae, p 1–44. *In* Howley PM, Knipe DM (ed), Fields Virology, 7 ed, vol 2: DNA Viruses. Lippincott Williams & Wilkins, Philadelphia, PA.

9. Wu Z, Graf FE, Hirsch HH. 2021. Antivirals against human polyomaviruses: Leaving no stone unturned. Rev Med Virol 31:e2220.

10. Leuzinger K, Kaur A, Wilhelm M, Frank K, Hillenbrand CA, Weissbach FH, Hirsch HH. 2022. Molecular Characterization of BK Polyomavirus Replication in Allogeneic Hematopoietic Cell Transplantation. J Infect Dis 227:888–900.

11. Leuzinger K, Kaur A, Wilhelm M, Hirsch HH. 2020. Variations in BK Polyomavirus Immunodominant Large Tumor Antigen-Specific 9mer CD8 T-Cell Epitopes Predict Altered HLA-Presentation and Immune Failure. Viruses 12:1476.

12. Wilhelm M, Kaur A, Wernli M, Hirsch HH. 2021. BK Polyomavirus-Specific CD8 T-Cell Expansion In Vitro Using 27mer Peptide Antigens for Developing Adoptive T-Cell Transfer and Vaccination. J Infect Dis 223:1410–1422.

13. Abend JR, Changala M, Sathe A, Casey F, Kistler A, Chandran S, Howard A, Wojciechowski D. 2017. Correlation of BK Virus Neutralizing Serostatus With the Incidence of BK Viremia in Kidney Transplant Recipients. Transplantation 101:1495–1505.

14. Pastrana DV, Ray U, Magaldi TG, Schowalter RM, Çuburu N, Buck CB. 2013. BK polyomavirus genotypes represent distinct serotypes with distinct entry tropism. J Virol 87:10105–10113.

15. Peretti A, Geoghegan EM, Pastrana DV, Smola S, Feld P, Sauter M, Lohse S, Ramesh M, Lim ES, Wang D, Borgogna C, FitzGerald PC, Bliskovsky V, Starrett GJ, Law EK, Harris RS, Killian JK, Zhu J, Pineda M, Meltzer PS, Boldorini R, Gariglio M, Buck CB. 2018. Characterization of BK Polyomaviruses from Kidney Transplant Recipients Suggests a Role for APOBEC3 in Driving In-Host Virus Evolution. Cell Host Microbe 23:628–635.e7.

16. McIlroy D, Hönemann M, Nguyen NK, Barbier P, Peltier C, Rodallec A, Halary F, Przyrowski E, Liebert U, Hourmant M, Bressollette-Bodin C. 2020. Persistent BK Polyomavirus Viruria is Associated with Accumulation of VP1 Mutations and Neutralization Escape. Viruses 12:824.

17. Bethge T, Hachemi HA, Manzetti J, Gosert R, Schaffner W, Hirsch HH. 2015. Sp1 sites in the noncoding control region of BK polyomavirus are key regulators of bidirectional viral early and late gene expression. J Virol 89:3396–3411.

18. Bethge T, Ajuh E, Hirsch HH. 2016. Imperfect Symmetry of Sp1 and Core Promoter Sequences Regulates Early and Late Virus Gene Expression of the Bidirectional BK Polyomavirus Noncoding Control Region. J Virol 90:10083–10101.

19. Gosert R, Rinaldo CH, Funk GA, Egli A, Ramos E, Drachenberg CB, Hirsch HH. 2008. Polyomavirus BK with rearranged noncoding control region emerge in vivo in renal transplant patients and increase viral replication and cytopathology. J Exp Med 205:841–852.

20. Martelli F, Giannecchini S. 2017. Polyomavirus microRNAs circulating in biological fluids during viral persistence. Rev Med Virol 27:e1927.

21. Jin L, Gibson PE, Booth JC, Clewley JP. 1993. Genomic typing of BK virus in clinical specimens by direct sequencing of polymerase chain reaction products. J Med Virol 41:11–17.

22. Jin L, Gibson PE, Knowles WA, Clewley JP. 1993. BK virus antigenic variants: sequence analysis within the capsid VP1 epitope. J Med Virol 39:50–56.

23. Cioni M, Leboeuf C, Comoli P, Ginevri F, Hirsch HH. 2016. Characterization of Immunodominant BK Polyomavirus 9mer Epitope T Cell Responses. Am J Transplant 16:1193–1206.

24. Leboeuf C, Wilk S, Achermann R, Binet I, Golshayan D, Hadaya K, Hirzel C, Hoffmann M, Huynh-Do U, Koller MT, Manuel O, Mueller NJ, Mueller TF, Schaub S, van Delden C, Weissbach FH, Hirsch HH, Swiss Transplant Cohort Study. 2017. BK Polyomavirus-Specific 9mer CD8 T Cell Responses Correlate With Clearance of BK Viremia in Kidney Transplant Recipients: First Report From the Swiss Transplant Cohort Study. Am J Transplant 17:2591–2600.

25. Kaur A, Wilhelm M, Wilk S, Hirsch HH. 2019. BK polyomavirus-specific antibody and T-cell responses in kidney transplantation: update. Curr Opin Infect Dis 32:575–583.

26. Sayers EW, Bolton EE, Brister JR, Canese K, Chan J, Comeau DC, Connor R, Funk K, Kelly C, Kim S, Madej T, Marchler-Bauer A, Lanczycki C, Lathrop S, Lu Z, Thibaud-Nissen F, Murphy T, Phan L, Skripchenko Y, Tse T, Wang J, Williams R, Trawick BW, Pruitt KD, Sherry ST. 2022. Database resources of the national center for biotechnology information. Nucleic Acids Res 50:D20–D26.

27. Altschul SF, Gish W, Miller W, Myers EW, Lipman DJ. 1990. Basic local alignment search tool. J Mol Biol 215:403–10.

28. Edgar RC. 2004. MUSCLE: multiple sequence alignment with high accuracy and high throughput. Nucleic Acids Res 32:1792–1797.

29. Meyder A, Nittinger E, Lange G, Klein R, Rarey M. 2017. Estimating Electron Density Support for Individual Atoms and Molecular Fragments in X-ray Structures. J Chem Inf Model 57:2437–2447.

30. Schwede T, Kopp J, Guex N, Peitsch MC. 2003. SWISS-MODEL: An automated protein homology-modeling server. Nucleic Acids Res 31:3381–3385.

31. Waterhouse A, Bertoni M, Bienert S, Studer G, Tauriello G, Gumienny R, Heer FT, de Beer TAP, Rempfer C, Bordoli L, Lepore R, Schwede T. 2018. SWISS-MODEL: homology modelling of protein structures and complexes. Nucleic Acids Res 46:W296–W303.

32. Benkert P, Biasini M, Schwede T. 2011. Toward the estimation of the absolute quality of individual protein structure models. Bioinformatics 27:343–350.

33. Schrödinger LLC, DeLano W. 2020. PyMOL, http://www.pymol.org/pymol.

34. Høie MH, Gade FS, Johansen JM, Würtzen C, Winther O, Nielsen M, Marcatili P. 2023. DiscoTope-3.0 - Improved B-cell epitope prediction using AlphaFold2 modeling and inverse folding latent representations. bioRxiv doi:10.1101/2023.02.05.527174:2023.02.05.527174.

35. Reynisson B, Alvarez B, Paul S, Peters B, Nielsen M. 2020. NetMHCpan-4.1 and NetMHCIIpan-4.0: improved predictions of MHC antigen presentation by concurrent motif deconvolution and integration of MS MHC eluted ligand data. Nucleic Acids Res 48:W449–W454.

36. Schymkowitz J, Borg J, Stricher F, Nys R, Rousseau F, Serrano L. 2005. The FoldX web server: an online force field. Nucleic Acids Res 33:W382–W388.

37. Salentin S, Schreiber S, Haupt VJ, Adasme MF, Schroeder M. 2015. PLIP: fully automated protein-ligand interaction profiler. Nucleic Acids Res 43:W443–W447.

38. Jumper J, Evans R, Pritzel A, Green T, Figurnov M, Ronneberger O, Tunyasuvunakool K, Bates R, Žídek A, Potapenko A, Bridgland A, Meyer C, Kohl SAA, Ballard AJ, Cowie A, Romera-Paredes B, Nikolov S, Jain R, Adler J, Back T, Petersen S, Reiman D, Clancy E, Zielinski M, Steinegger M, Pacholska M, Berghammer T, Bodenstein S, Silver D, Vinyals O, Senior AW, Kavukcuoglu K, Kohli P, Hassabis D. 2021. Highly accurate protein structure prediction with AlphaFold. Nature 596:583–589.

39. Lilyestrom W, Klein MG, Zhang R, Joachimiak A, Chen XS. 2006. Crystal structure of SV40 large T-antigen bound to p53: interplay between a viral oncoprotein and a cellular tumor suppressor. Genes Dev 20:2373–2382.

40. Cuesta I, Nunez-Ramirez R, Scheres SH, Gai D, Chen XS, Fanning E, Carazo JM. 2010. Conformational rearrangements of SV40 large T antigen during early replication events. J Mol Biol 397:1276–86.

41. Loeber G, Parsons R, Tegtmeyer P. 1989. The zinc finger region of simian virus 40 large T antigen. J Virol 63:94–100.

42. Chang YP, Xu M, Machado ACD, Yu XJ, Rohs R, Chen XS. 2013. Mechanism of origin DNA recognition and assembly of an initiator-helicase complex by SV40 large tumor antigen. Cell Rep 3:1117–1127.

43. Marrama D, Mahita J, Sette A, Peters B. 2022. Lack of evidence of significant homology of SARS-CoV-2 spike sequences to myocarditis-associated antigens. EBioMedicine 75:103807.

44. Amorim AR, Mendes GS, Santos N. 2022. Genotyping of human polyomavirus 1 detected in saliva. Gene Reports 27:101629.

45. Sorin MN, Di Maio A, Silva LM, Ebert D, Delannoy CP, Nguyen NK, Guerardel Y, Chai W, Halary F, Renaudin-Autain K, Liu Y, Bressollette-Bodin C, Stehle T, McIlroy D. 2023. Structural and functional analysis of natural capsid variants suggests sialic acid-independent entry of BK polyomavirus. Cell Rep 42:112114.

46. Neu U, Allen SA, Blaum BS, Liu Y, Frank M, Palma AS, Ströh LJ, Feizi T, Peters T, Atwood WJ, Stehle T. 2013. A structure-guided mutation in the major capsid protein retargets BK polyomavirus. PLoS Pathog 9:e1003688.

47. Chen XS, Stehle T, Harrison SC. 1998. Interaction of polyomavirus internal protein VP2 with the major capsid protein VP1 and implications for participation of VP2 in viral entry. EMBO J 17:3233–3240.

48. Willhelm M, Wilk S, Kaur A, Hirsch HH, Swiss Transplant Cohort Study. 2019. Can HLA-B51 Protect Against BKPyV-DNAemia? Transplantation 103:e384–e385.

49. Solis M, Velay A, Porcher R, Domingo-Calap P, Soulier E, Joly M, Meddeb M, Kack-Kack W, Moulin B, Bahram S, Stoll-Keller F, Barth H, Caillard S, Fafi-Kremer S. 2018. Neutralizing Antibody-Mediated Response and Risk of BK Virus-Associated Nephropathy. J Am Soc Nephrol 29:326–334.

50. Pastrana DV, Brennan DC, Cuburu N, Storch GA, Viscidi RP, Randhawa PS, Buck CB. 2012. Neutralization serotyping of BK polyomavirus infection in kidney transplant recipients. PLoS Pathog 8:e1002650.

51. Domingo-Calap P, Schubert B, Joly M, Solis M, Untrau M, Carapito R, Georgel P, Caillard S, Fafi-Kremer S, Paul N, Kohlbacher O, González-Candelas F, Bahram S. 2018. An unusually high substitution rate in transplant-associated BK polyomavirus in vivo is further concentrated in HLA-C-bound viral peptides. PLoS Pathog 14:e1007368.

52. Stavrou S, Ross SR. 2015. APOBEC3 Proteins in Viral Immunity. J Immunol 195:4565–4570.

53. Harris RS, Dudley JP. 2015. APOBECs and virus restriction. Virology 479–480:131-145.

54. Wunderink HF, Haasnoot GW, de Brouwer CS, van Zwet EW, Kroes ACM, de Fijter JW, Rotmans JI, Claas FHJ, Feltkamp MCW. 2019. Reduced Risk of BK Polyomavirus Infection in HLA-B51-positive Kidney Transplant Recipients. Transplantation 103:604–612.

