## Supplementary figures and images for "Structural implications of BK polyomavirus sequence variations in the major viral capsid protein Vp1 and large T-antigen: a computational study"

### Supplemental Figure 1

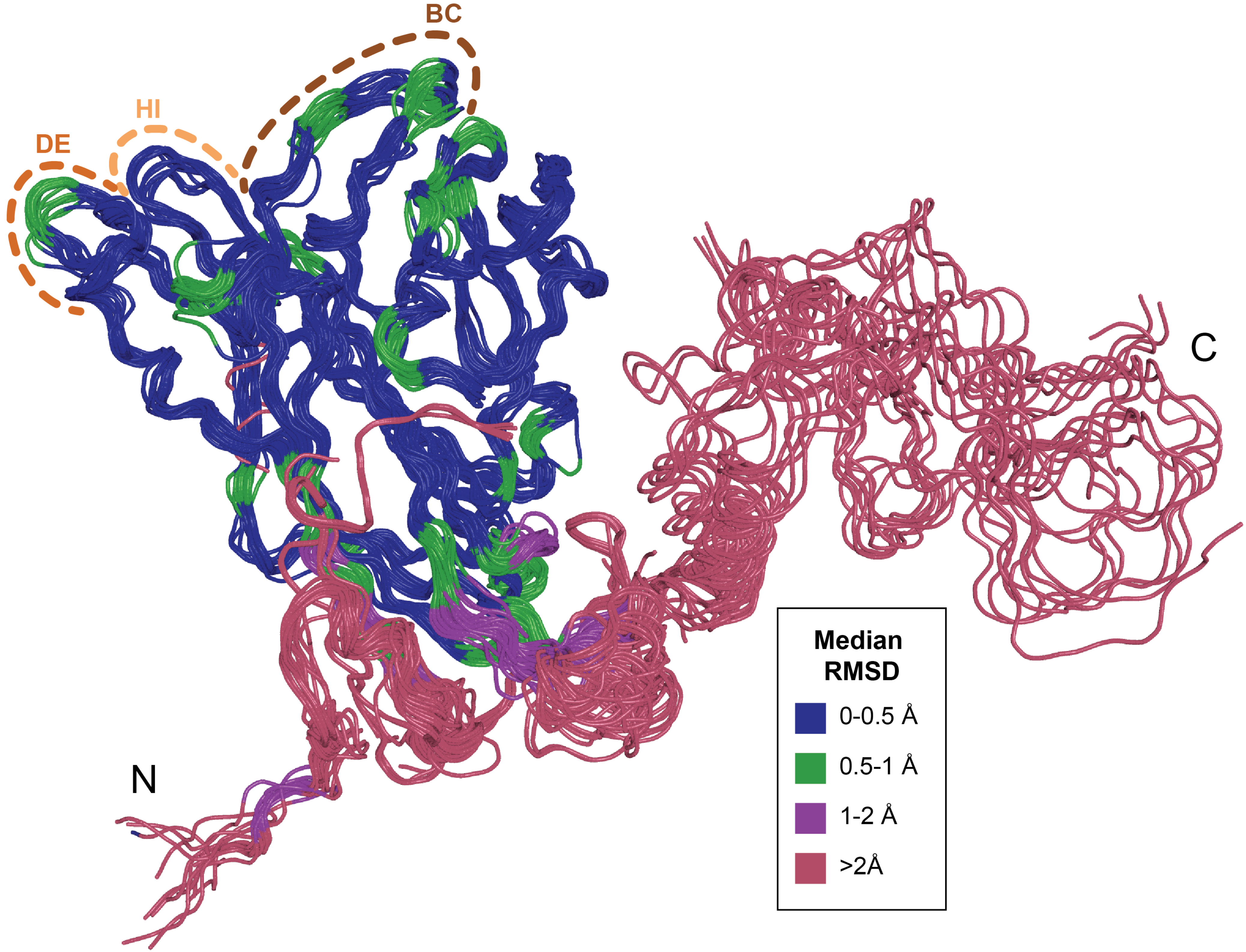
